# Spatiotemporal dynamics of adoptively transferred stem-like CD8^+^ T cells in the tumor microenvironment following vaccination

**DOI:** 10.64898/2026.05.12.724323

**Authors:** Dalton Hermans, Sloane C Fussell, Andrei Ramirez-Valdez, Sam Shepard, Remi Poulard, Sonia Zumalave, Benjamin Sievers, Christopher M Garliss, Vincent L Coble, Geoffrey M Lynn, Andrew S Ishizuka, Isidro Cortes-Ciriano, Robert A Seder

## Abstract

Adoptive cell therapy (ACT) of tumor-specific T cells can improve survival in a subset of cancer patients. Current ACT approaches may be limited by using highly differentiated T cells which can be inhibited by an immunosuppressive tumor microenvironment (TME). Here, we developed an approach to optimize ACT and used spatial transcriptomics to show how stem-like and effector CD8^+^ T cells differentially mediate tumor control following vaccination. Spatial transcriptomic profiling of the TME showed that ACT with stem-like T cells followed by intravenous vaccination prevented immune exclusion, increased infiltration of pro-inflammatory macrophages, and reprogrammed tumor cells to upregulate Type I and Type II IFN signaling and apoptotic gene programs. The protective transcriptomic signature of the TME in this ACT model contained overlapping biomarkers with patients who responded to ACT therapy. This approach demonstrates synergy between transferred stem-like T cells and intravenous vaccination to transcriptionally remodel the TME and enhance tumor control.

## Introduction

Adoptive cell therapy (ACT) has become a transformative modality in cancer treatment, particularly for patients with refractory hematologic malignancies and select solid tumors^1^. Current cell therapies are largely divisible into three main approaches: tumor-infiltrating lymphocyte (TIL) therapy, chimeric antigen receptor T cell (CAR-T) therapy, and T cell receptor (TCR-T) platforms. Cell therapies are multi-step procedures whose efficacy is determined by three stages^2^: 1) expansion of T cells following *in vitro* activation; 2) activation and expansion of transferred T cells *in vivo*; and 3) infiltration and anti-tumor function of T cells in the tumor microenvironment (TME). Each of these steps has limitations that effect tumor control. Thus, a broad-based approach to optimize ACT immunotherapy at each stage of the process is needed to improve clinical efficacy.

A major advantage of using ACT for tumor immunotherapy is the ability to transfer large numbers of T cells, which may be required for individuals with substantial tumor burden or metastatic disease^3^. In addition to magnitude, the quality of the T cell can also have an important role for optimizing tumor control. Indeed, T cells in less-differentiated activation states have a greater anti-tumor efficacy than differentiated effector cells (T_eff_) following adoptive transfer^4,5^. In particular, stem-cell memory T cells (T_scm_) with enhanced capacity for self-renewal, proliferation, and differentiation into effector cells have superior anti-tumor effects and enhanced responsiveness to checkpoint inhibition therapy (CPI)^6–9^. A key goal for ACT is to allow for expansion of T cells during the *in vitro* activation phase while maintaining a less differentiated pool of stem-like cells. Genetic manipulation, cytokine stimulation, and small-molecule inhibitors have been previously used to maintain T_scm_ cells in culture^10–13^. Indeed, CAL-101 (Idelalisib), a small-molecule inhibitor of the delta isoform of phosphatidylinositol 3-kinase (PI3k-δ)^14^ has been shown to restrict differentiation during *in vitro* T cell stimulation leading to maintenance of stem-like T cells^15–18^, thereby providing a simple approach to optimize the magnitude and quality of T cells following *in vitro* expansion.

An additional consideration for optimizing ACT is to improve T cell function in the TME^19^. Analysis of the TME by various technologies has revealed extensive heterogeneity across patients that are associated with variable responsiveness to immune-based therapy^20–22^. Several suppressive mechanisms have been elucidated by which tumor cells evade detection and elimination by the immune system, including immunoediting, immune exclusion, production of soluble suppressive factors (e.g. TGF-β, IL-10), and recruitment of anti-inflammatory and/or suppressive immune cells^23–25^. Thus, therapies like ACT or therapeutic vaccines focusing on enhancing T cell immunity may be improved from combination with TME-modifying agents^26^. We previously reported^27,28^ in various mouse tumor models that neo-antigen vaccines in the therapeutic setting can boost tumor-specific T cells and induce systemic type I IFN, leading to TME remodeling and effective tumor control. In humans, therapeutic cancer vaccine trials, particularly in the adjuvant setting (following surgical resection) have been shown to prevent recurrent disease^29–32^. However, clinical studies assessing the effect of therapeutic vaccines on more established and metastatic tumors have been less successful^33–35^. This may be due to an insufficient magnitude of T cells required to regress advanced or metastatic solid tumors or the inability to function in an immunosuppressive TME^26^.

In this report, we hypothesized that a comprehensive approach which increases the magnitude of stem-like T cells for ACT combined with the immunomodulatory benefits of intravenous (IV) vaccination to expand T cells *in vivo* and modulate the TME would be optimal for tumor control. An additional focus was to visualize how stem-like T cells were distributed in the TME and identify transcriptomic signatures of protection *in situ* to elucidate potential protective mechanisms and define biomarkers. While single-cell RNA sequencing (scRNAseq) and multiplexed microscopy methods have been used to define the immune cell composition and activity in the TME^36–38^, these analysis are limited by an absence of the spatial context and sufficient markers to perform unbiased analysis, respectively. Here, to evaluate the effect of ACT followed by vaccination in the TME, we performed unbiased, subcellular spatial transcriptomics to define the *in situ* spatial dynamics between tumor and immune cells. Last, we found that biomarkers derived from the TME following treatment with T_scm_ cells and vaccination were correlated with genes upregulated in human responders to ACT and a pan-cancer cohort of patients with improved survival outcomes. This approach for optimizing ACT focused on remodeling the TME may guide future therapeutic approaches.

## Results

### Magnitude and quality of CD8^+^ T cells following *in vitro* activation with CAL-101

The first step toward optimizing ACT is to expand the magnitude of cells during *in vitro* culture while maintaining specific phenotypic and transcriptional stem-like features for improved *in vivo* function. To model this, naïve OT-1 CD8^+^ T cells were activated in the presence of either 2.5μM CAL-101, a small-molecule inhibitor^15,16^ of the delta isoform of phosphatidylinositol 3-kinase (PI3k-δ), or control DMSO (**Figure 1A**). After four days of stimulation, activated CD8^+^ T cells were assessed using PD1 and TCF1, canonical markers for characterizing stem-like (PD1^+^TCF1^+^) and more differentiated effector (PD1^+^TCF1^-^) cells. Naïve OT-1 cells cultured with CAL-101 were 80.4% PD1^+^TCF1^+^ which will be designated hereafter as stem-cell memory T cells (T_scm_). Conversely, cells cultured in the absence of CAL-101 were 67.5% PD1^+^TCF1^-^, and will be designated as effector T cells (T_eff_) (**Figure 1B**). Within the PD1^+^TCF1^+^ populations, there was also significantly higher mean fluorescence intensity (MFI) of Slamf6, an additional marker used to define functional stem-like cells^39^ in the T_scm_ cells compared to the T_eff_ cells (**Figures 1C and 1D**). As CAL-101 increased the proportion of PD1^+^TCF1^+^ cells compared to the control culture (**Figure 1E**), phenotypic markers of memory T cells were also assessed. 65% of CD8^+^ T cells cultured with CAL-101 expressed the lymph-node homing molecule CD62L, a marker of central memory cells (**Figure 1F**) compared to 31.6% in the T_eff_ cells. Importantly, while inhibition of PI3k by CAL-101 preserved the stem-like quality and maintained a greater proportion of central memory cells, the magnitude of expansion was retained (**Figure 1G**).

**Figure 1.**
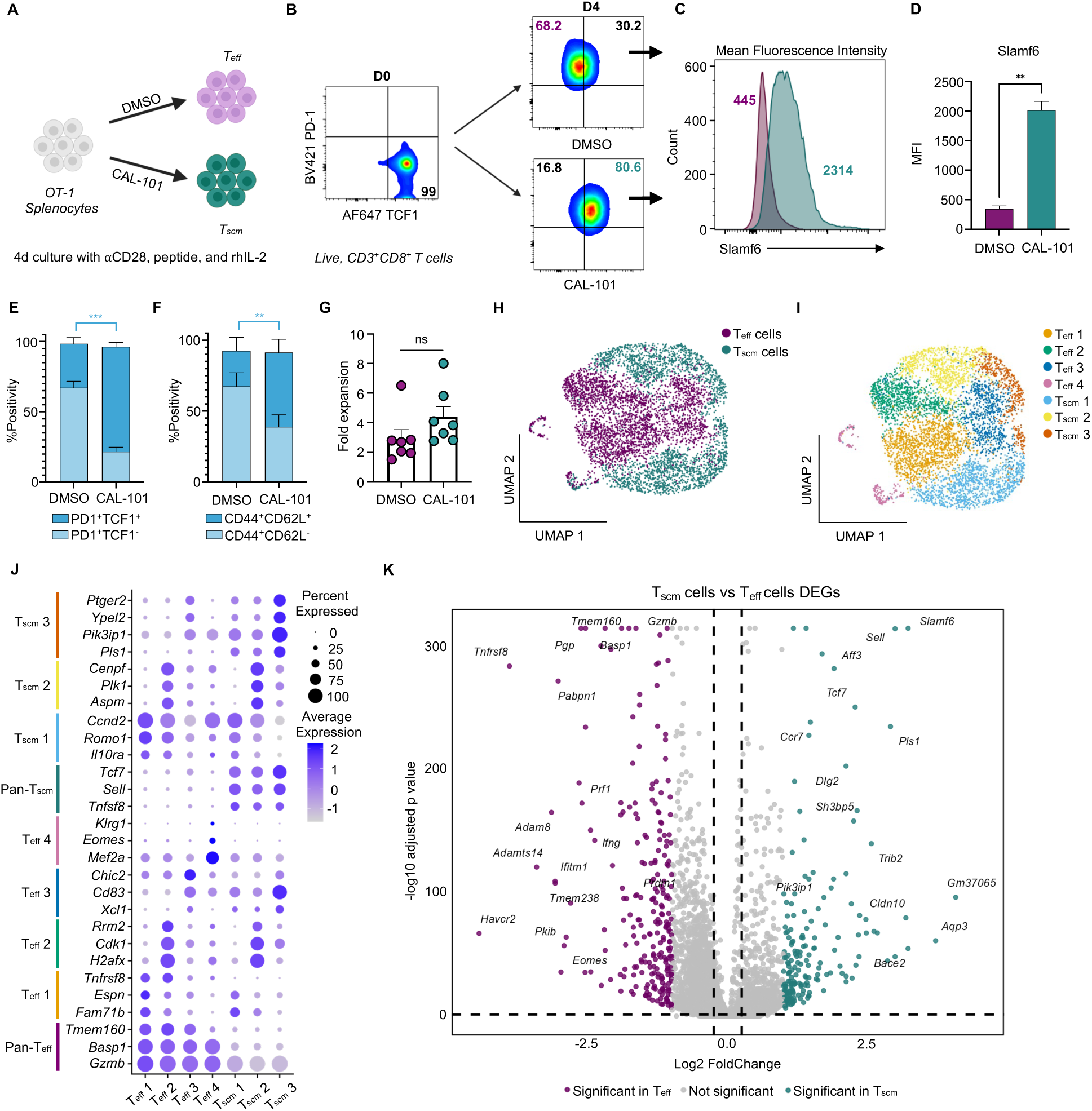
Magnitude and quality of CD8+ T cells following in vitro activation with CAL-101 (A) Schematic of cell culture protocol. (B) Flow cytometry of PD1vsTCF1 expression gated on live, CD3^+^CD8^+^ T cells on Day 0 before start of culture and CD8^+^ T cells after four day culture with DMSO or CAL-101. (C) Histogram of Slamf6 MFI from the PD1^+^TCF1^+^ cell populations in DMSO (purple) or CAL-101 (green) treated cells. (D) Bar graph quantitating MFI of Slamf6 in PD1^+^TCF1^+^ DMSO (purple) or CAL-101 (green) treated cells (n = 3). Statistics accessed by paired, two-tailed T test. (E) Bar graph showing the percentages of PD1 and TCF1 subpopulations in DMSO or CAL-101 treated cells (n=7). Statistics accessed by paired, two-tailed T test. (F) Bar graph showing the percentages of CD44 and CD62L subpopulations in DMSO or CAL-101 treated cells (n=6). Statistics accessed by paired, two-tailed T test. (G) Bar graph indicating the fold-expansion of CD8^+^ T cells following the four day culture period. (H) UMAP of scRNAseq data showing Teff cells (purple) and Tscm cells (green). (I) UMAP of scRNAseq data showing clusters composed of Teff or Tscm cells. (J) Dot plot showing average and percent expression of key DEGs from each cluster. (K) Volcano plot showing significant (p < 0.05) DEGs that are upregulated (green) or downregulated (purple) by comparing Tscm cells with Teff cells.

To further analyze the cultured cell products, scRNAseq was performed on CD8^+^ T cells that were activated and treated with either CAL-101 or DMSO. Following dimensional reduction, cells clustered based on treatment group with four discrete T_eff_ clusters and three discrete T_scm_ clusters (**Figures 1H and 1I**). Compared to T_scm_ cells, T_eff_ cells were enriched for Hallmark gene sets that included TNFα signaling via NFκB, PI3k/mTOR signaling, hypoxia, and MYC targets (**Figures S1A and S1B**). Unbiased differential gene expression analysis showed that T_eff_ cells were highly differentiated by the expression of *Gzmb*, *Tnfrsf8*, *Tmem160*, and *Basp1* while T_scm_ cells expressed *Slamf6*, *Tcf7*, *Sell*, and *Pls1* (**Figures 1J, S1C, and S1D**). The analysis also revealed that T_eff_ cells significantly upregulated traditional effector genes *Havcr2, Eomes, and Ifng* while T_scm_ cells upregulated *Slamf6 and Ccr7* (**Figure 1K**). Clusters in both the T_eff_ and T_scm_ cohorts displayed a range of activation states that suggest subtle differences in amount of TCR stimulation^40^ and CAL-101 drug inhibition. Notably, the most stem-like cluster T_scm_ 3 is marked by its differential expression of *Pik3ip1*, a negative regulator of PI3k, suggesting that cells receiving the highest amount of PI3k inhibition have the greatest preservation of stemness (**Figure 1J**). T_scm_ 3 also differentially expresses the cellular senescence gene *Ypel2*^41^ and the EP_2_ receptor *Ptger2*^42^ which may restrict IL-2-mediated differentiation signals during the culture. The heterogeneity within the T_eff_ cells ranged from a highly differentiated T_eff_ 4 cluster enriched for *Klrg1* and *Eomes* and a less differentiated T_eff_ 3 cluster containing the largest expression amongst T_eff_ cells of *Tcf7*, *Ptger2*, *Ypel2*, and *Pik3ip1*. T_eff_ 3 also expresses *Xcl1*, a chemokine previously reported^43,44^ in stem-like T cells, at similar levels as the T_scm_ clusters (**Figure 1J**). Together these data show two distinct transcriptional and phenotypic types of CD8^+^ T cells for ACT depending on the presence of CAL-101 during culture.

### T cell expansion and tumor control following ACT and vaccination

To assess the *in vivo* functionality of T_scm_ and T_eff_ cells, cells were adoptively transferred into mice that had received an ovalbumin-expressing B16 melanoma tumor (B16-Ova). Tumors were inoculated on the flank of CD45.1^+^ mice and allowed to grow for 12 days reaching ∼100mm^3^ before transferring 10^5^ CD45.2^+^ T_scm_ or T_eff_ cells. We used the SNAPVax^TM^ vaccine^45^, a vaccine platform consisting of self-assembling nanoparticles that codeliver peptide antigens and toll-like receptor -7 and -8 agonists (hereafter referred to as SNP-7/8a) to enhance transferred cells. The therapeutic effect of T_scm_ and T_eff_ cells with or without vaccination (Vax) with Ova:SNP-7/8a three days after transfer was assessed (**Figure 2A**). Mice that received no ACT or vaccination alone served as negative controls. Mice that received ACT with T_eff_ or T_scm_ cells alone showed a similar delay in tumor growth compared to mice receiving no cells or therapeutic vaccination alone (**Figure 2B**); however, mice that received T_scm_ cells + Vax had significantly improved anti-tumor control and reduction in mortality compared to ACT with T_eff_ cells + Vax (**Figure 2B**).

**Figure 2.**
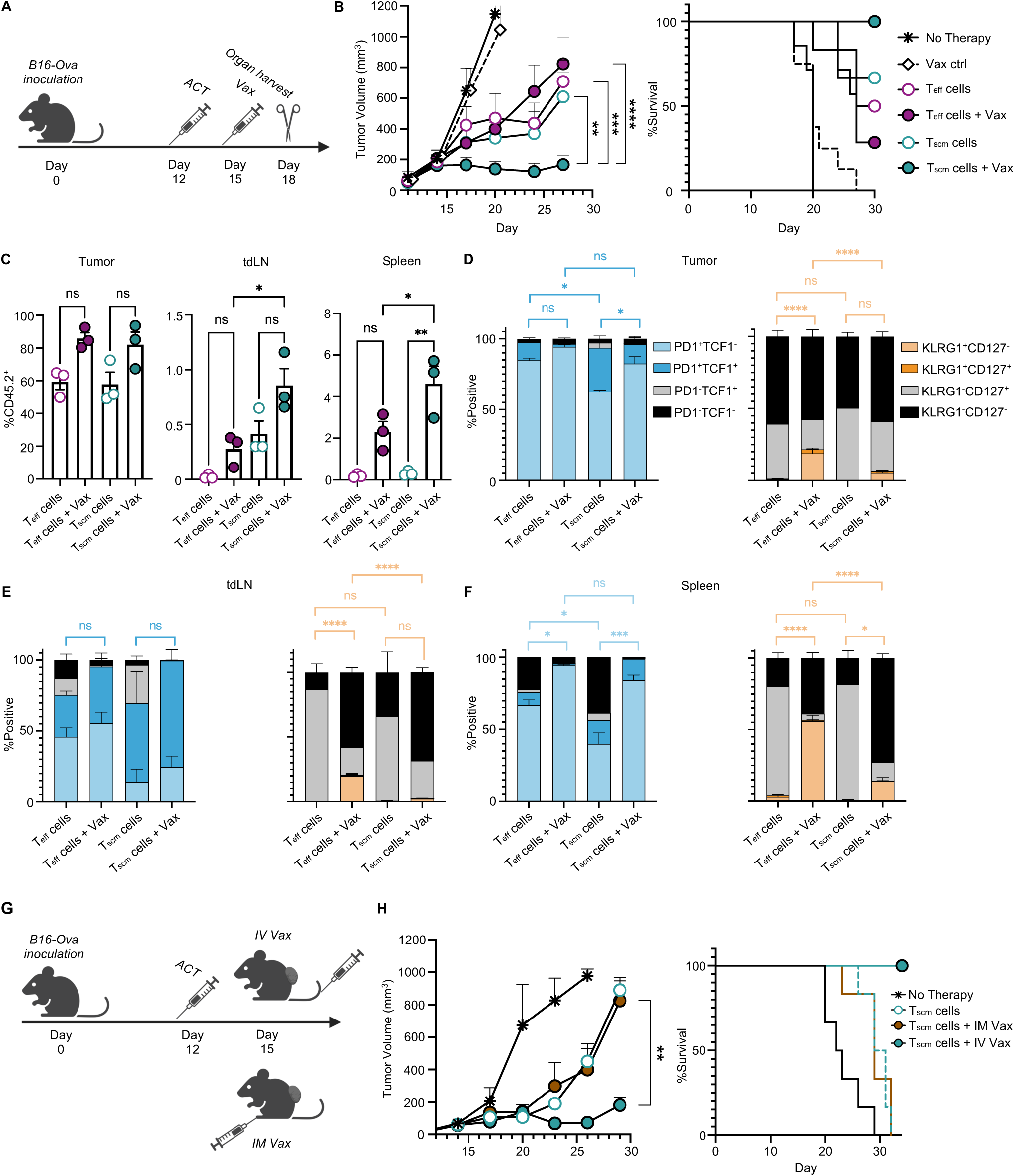
T cell expansion and tumor control following ACT and vaccination. (A) Experimental schematic of mouse model testing the effect of ACT and vaccination. (B) Tumor curve (left) following treatment of Teff cells or Tscm cells with or without vaccination (n=7) and (right) survival curves. Tumor statistics accessed by two-way ANOVA. (C) Bar graphs showing the frequency of CD45.2^+^ transferred cells in the tumor, tdLN, and spleen by treatment group (n=3). Statistics accessed by ANOVA. (D) Stacked bar graphs indicating the percentages of PD1 and TCF1 (left) or KLRG1 and CD127 (right) subpopulations in tumor (n=3). Statistics assessed by ANOVA. (E) Stacked bar graphs indicating the percentages of PD1 and TCF1 (left) or KLRG1 and CD127 (right) subpopulations in tdLN (n=3). Statistics assessed by ANOVA. (F) Stacked bar graphs indicating the percentages of PD1 and TCF1 (left) or KLRG1 and CD127 (right) subpopulations in spleen (n=3). Statistics assessed by ANOVA. (G) Experimental schematic of mouse model testing the effect of vaccine route following ACT. (H) Tumor curve (left) following treatment of Tscm cells with IV or IM vaccination (n=6) and (right) survival curves. Tumor statistics accessed by two-way ANOVA.

As vaccination is required for optimal protection following ACT with T_scm_ cells, we assessed T cell responses of transferred CD45.2^+^ cells in the tumor, tumor-draining lymph node (tdLN), and spleen from animals three days after vaccination (**Figure 2C**). In the tumor tissue, there was a similar increase in the number of CD45.2^+^ T_eff_ and T_scm_ cells following vaccination compared to the cells alone (**Figure 2C**). For the tdLN and spleen, there was an ∼2-fold increase of transferred CD45.2^+^ T cells in in the T_scm_ cells + Vax group compared to the T_eff_ cells + Vax group (**Figure 2C**). The expression of PD1, TCF1, CD127, and KLRG1, canonical markers defining the differentiate state of memory and effector T cells^46^, was assessed. The phenotype of transferred cells varied based on organ and treatment group (**Figures 2D-2F and S2A**). T_scm_ cells in tumor had a higher proportion of PD1^+^TCF1^+^ cells than T_eff_ cells (**Figure 2D**). CD45.2^+^ T cells from T_eff_ cells + Vax group were >90% PD1^+^TCF1^-^ and had the largest population (∼19%) of KLRG1^+^CD127^-^ cells, which define highly differentiated effector T cells (**Figure 2D**). In the tdLN, there was a high proportion of TCF1 expression in CD45.2^+^ T_scm_ cells with or without vaccination (75 and 83%, respectively), while only ∼40% of T_eff_ cells with or without vaccination were TCF1^+^ (**Figure 2E**). Vaccination did not significantly alter the KLRG1 expression of T_scm_ cells recovered from the tdLN, however, similar to tumor tissue, vaccination converted a subset (∼20%) of T_eff_ cells towards a KLRG1^+^CD127^-^ phenotype (**Figure 2E**). Vaccine induced conversion towards a KLRG1^+^CD127^-^ phenotype was strongest in the spleen, where ∼56% of splenic CD45.2^+^ T_eff_ cells + Vax were KLRG1^+^CD127^-^ compared to 14% of T_scm_ cells + Vax (**Figure 2F**). Together, these data show that mice treated with T_scm_ cells + Vax contain a greater proportion of stem-like TCF1^+^ T cells and a lower frequency of KLRG1 in the tdLN and spleen compared to mice treated with T_eff_ cells + Vax (**Figures 2D-2F**), suggesting maintenance of less differentiated cells *in vivo*.

To substantiate the protective effect of this ACT and vaccination approach in a separate tumor model, an MC38 colon carcinoma tumor line expressing a KVP-mutated gp100 protein^47^ was assessed (**Figure S2B**). Naïve CD8^+^ p-mel T cells specific for gp100 were activated *in vitro* in the presence or absence of CAL-101 and used for ACT. MC38-gp100 tumors were implanted for 14 days before being treated with 10^6^ T_eff_ or T_scm_ cells, with or without vaccination with gp100:SNP-7/8a. In this model, treatment groups (including T_eff_ and T_scm_ cells alone) achieved complete tumor clearance in most animals and long-term survival (**Figure S2C and S2D**). To create a more aggressive tumor model, MC38-gp100 tumor cells were mixed^48^ with wild-type MC38 tumor cells in a 50:50 ratio (**Figure S2E**). Remarkably, removing the target gp100 antigen from 50% of the tumor cells resulted in significant differences in the efficacy of the treatment groups. Similar to the B16-OVA melanoma model, ACT with T_scm_ or T_eff_ cells alone was unable to control the mixed tumor. Vaccination also had no therapeutic effect on transferred T_eff_ cells for delay of tumor growth or overall survival (**Figure S2F and S2G**). By contrast, animals that received T_scm_ cells followed by vaccination had significant control of tumor and increased survival (**Figure S2F and S2G**). These data confirm in a second model that ACT with T_scm_ cells followed by intravenous vaccination was most effective for controlling tumor.

### Effect of vaccine route on immunity and protection following ACT

A potentially critical aspect of tumor control following ACT may be the requirement for intravenous vaccination, which has been shown to enhance immunogenicity and protection against tumors compared to conventional routes of immunization^27,28,49^. Indeed, IV delivery with the SNP-7/8 vaccine in a therapeutic tumor model using neoantigens was required to control tumor growth compared to subcutaneous (SC) or intramuscular (IM) delivery^28^. Thus, to determine if vaccine route was also a key determinant in this ACT model in which T cells are already present at high frequency *in vivo*, we tested whether intramuscular vaccination would mediate control of tumor (**Figure 2G**). In contrast to protection by ACT with T_scm_ cells + IV vaccination, mice treated with T_scm_ cells + IM vaccination did not have significantly different tumor protection or survival from mice receiving T_scm_ cells alone (**Figure 2H**). Consistent with this lack of tumor control, IM vaccination failed to expand transferred T_scm_ cells in blood while IV vaccination significantly increased the transferred T_scm_ cells to ∼20% (**Figure S2H**). These findings are consistent with previous data^27^ showing that the SNP-7/8a vaccine is retained at the injection site or local draining lymph node following IM injection, while IV administered SNP-7/8a traffics to the spleen, tumor, and tdLN. The vaccine-induced expansion of transferred T_scm_ cells was also associated with increased proportions of PD1^+^TCF1^-^ effector T cells (**Figure S2I**) and a higher proportion of CD127^-^ cells (**Figure S2J**), consistent with activation *in vivo*.

### Requirement of antigen and innate activation by IV vaccination for tumor control

The SNP-7/8a vaccine provides simultaneous delivery of the selected antigen with innate stimulation mediated by a toll-like receptor-7/8 agonist^45^, allowing for T cell activation, expansion, and induction of systemic innate immunity. Here, to determine the contribution of antigen, innate stimulation, or both on immunogenicity and tumor control by the SNP-7/8a vaccine following ACT we compared three separate vaccine formulations: 1) Ova:SNP, composed of Ova peptide without innate stimulation and testing the effect of antigen alone; 2) E7:SNP-7/8a, composed of an irrelevant E7 peptide linked to TLR-7/8a and testing the effect of innate stimulation alone; and 3) Ova:SNP-7/8a, composed of Ova peptide linked to TLR-7/8a and testing the effect of both antigen and innate stimulation (**Figure S3A**). ACT with T_scm_ followed by vaccination with antigen alone (T_scm_ cells + Ova:SNP) had significantly decreased tumor control compared to the T_scm_ cells + Ova:SNP-7/8a (**Figure 3A**). ACT with innate stimulation alone (T_scm_ cells + E7:SNP-7/8a) showed enhanced tumor control that was not significantly different from mice treated with T_scm_ cells + Ova:SNP-7/8a, however, the latter did have improved overall survival (**Figure 3A and 3B**). There was no significant differences in tumor control or survival for mice treated with T_eff_ cells and vaccinated with Ova:SNP-7/8 or E7:SNP-7/8a (**Figure S3B**). However, mice treated with T_eff_ cells + Ova:SNP had decreased survival compared to mice treated with T_eff_ cells + Ova:SNP-7/8a (**Figure S3C**).

**Figure 3.**
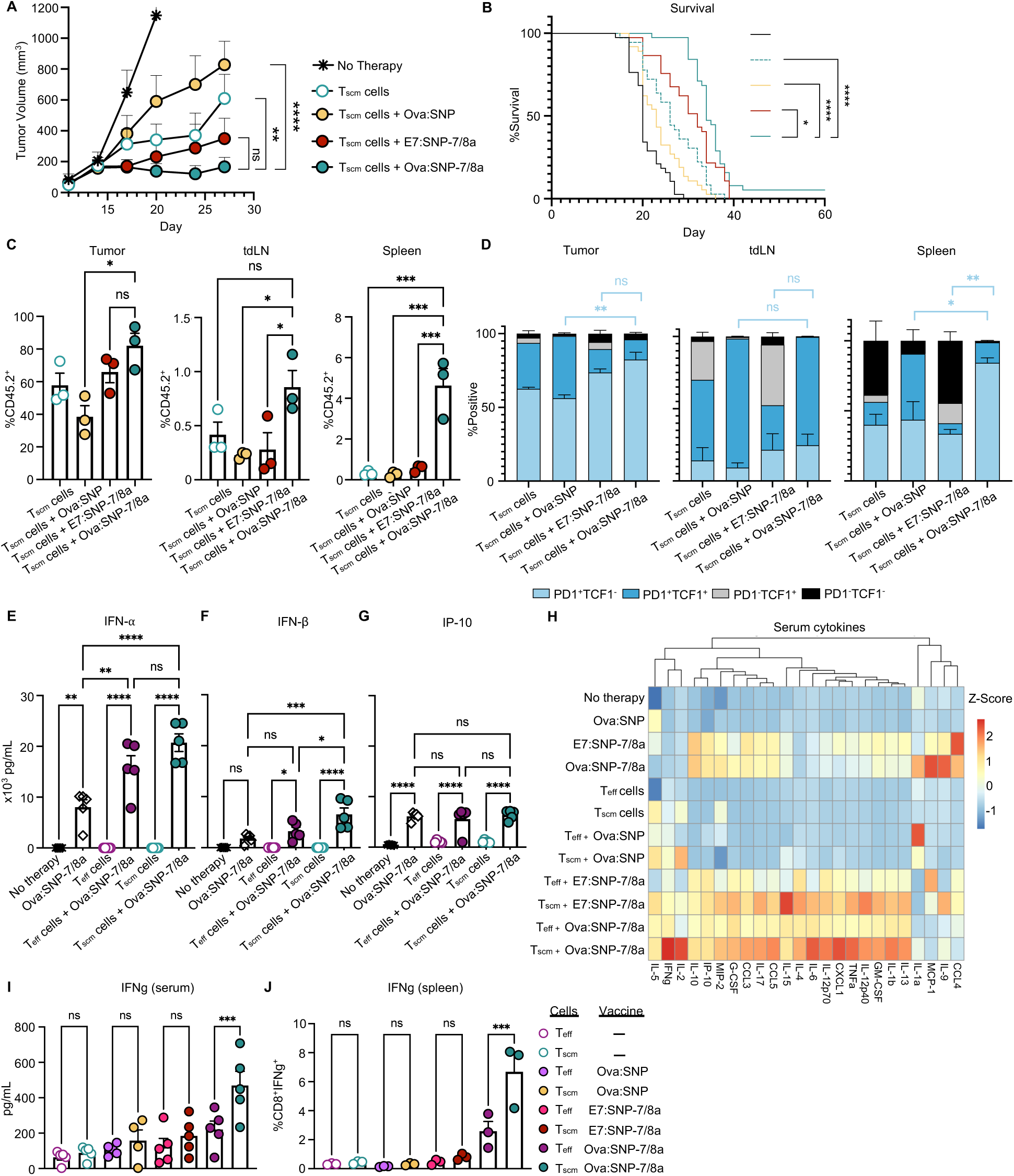
Requirement of antigen and innate activation by IV vaccination for tumor control. (A) Tumor curve following treatment of Tscm cells with various vaccine conjugates (n=7). Statistics accessed by two-way ANOVA. (B) Survival curve following treatment of Tscm cells with various vaccine conjugates (n=35). Statistics accessed by log-rank test. (C) Bar graphs showing the frequency of CD45.2^+^ transferred cells in the tumor, tdLN, and spleen by treatment group (n=3). Statistics accessed by ANOVA. (D) Stacked bar graphs indicating the percentages of PD1 and TCF1 subpopulations in tumor, tdLN, and spleen (n=3). Statistics assessed by ANOVA. (E) Bar graph showing the serum cytokine levels (pg/mL) of IFN-α 6 hours post-vaccination in indicated treatment groups (n=5). Statistics assessed by ANOVA. (F) Bar graph showing the serum cytokine levels (pg/mL) of IFN-β 6 hours post-vaccination in indicated treatment groups (n=5). Statistics assessed by ANOVA. (G) Bar graph showing the serum cytokine levels (pg/mL) of IP-10 6 hours post-vaccination in indicated treatment groups (n=5). Statistics assessed by ANOVA. (H) Heatmap of Z-scored serum cytokine data taken 6 hours post-vaccination in indicated treatment groups (n=5). (I) Bar graph showing the serum cytokine levels (pg/mL) of IFN-γ 6 hours post-vaccination in indicated treatment groups (n=5). Statistics assessed by ANOVA. (J) Bar graph showing frequency by flow cytometry of IFN-γ^+^ cells among live, CD8^+^ T cells in the spleen three days following vaccination (n=3). Statistics assessed by ANOVA.

Immune analysis of transferred CD45.2^+^ cells in the tumor, tdLN, and spleen revealed differences in responses to the different vaccine formulations. Ova:SNP lowered the proportion of T_scm_ cells in the tumor, while Ova:SNP-7/8a significantly increased the proportion of T_scm_ cells in the tdLN and spleen (**Figure 3C**). T_eff_ cells showed similar trends in the tumor, tdLN, and spleen when tested against the different vaccine formulations, albeit at lower magnitudes (**Figure S3D**). Different vaccine conjugates elicited distinct phenotypes in transferred T cells, with T_scm_ cells receiving Ova:SNP showing a similar phenotype as T_scm_ cells alone (**Figure 3D**). Innate stimulation with or without antigen produced T cells with less TCF1 expression. Of note, in the spleen there was a significantly larger population of PD1^+^TCF1^-^ in the group that received T_scm_ cells + Ova:SNP-7/8a compared to T_scm_ cells + E7:SNP-7/8a (**Figure 3D**). Vaccination with Ova:SNP-7/8a also induced a greater proportion of PD1^+^TCF1^-^ cells in T_eff_ cells compared to E7:SNP-7/8a (**Figure S3E**). Together, these data show that antigen alone is not sufficient or may even be detrimental for enhancing T cell function while innate stimulation alone may provide some benefit following ACT with T_scm_ cells (**Figure 2B**). Optimal therapy is achieved when ACT is combined with a vaccine containing both target antigen and innate stimulation.

### Innate activation *in vivo* following ACT with T_eff_ and T_scm_ cells and vaccination

To assess how the SNP vaccine altered innate activation *in vivo* and determine whether ACT with T_eff_ or T_scm_ cells affects these responses, we assessed Type I IFN and IP-10, two canonical innate cytokines induced by TLR-7/8, six hours following vaccination. Mice that received only an IV vaccination with Ova:SNP-7/8a or E7:SNP-7/8a had comparable and increased production of IFN-α, IFN-β, and IP-10 compared to mice that received vaccination with antigen alone (Ova:SNP) or no therapy (**Figures S3F-S3H**). ACT with T_scm_ cells or T_eff_ cells followed by IV vaccination with Ova:SNP-7/8a had significantly increased serum levels of IFN-α compared to mice that received vaccination alone (**Figure 3E**). Serum levels of IFN-β were significantly higher following ACT with T_scm_ cells compared to T_eff_ cells after vaccination compared to vaccination alone (**Figure 3F).** Serum levels of IP-10 were similar in mice that received T_scm_ cells or T_eff_ cells followed by Ova:SNP-7/8a compared to Ova:SNP-7/8a vaccination alone (**Figure 3G**). In mice treated with T_scm_ cells, there was a significant increase in IFN-α and IFN-β following vaccination with Ova:SNP-7/8a compared with E7:SNP-7/8a (**Figure S3F and S3G**). However, in mice treated with T_eff_ cells, similar levels of IFN-α and IFN-β were observed following vaccination with Ova:SNP-7/8a or E7:SNP-7/8a (**Figure S3F and S3G**). By contrast, mice treated with T_eff_ cells had similar levels of IFN-α and IFN-β following vaccination with Ova:SNP-7/8a or E7:SNP-7/8a (**Figure S3F and S3G**). In contrast to the effect of ACT for increasing IFN-α and IFN-β following vaccination, IP-10 levels were similar in mice that received Ova:SNP-7/8a or E7:SNP-7/8a, with or without ACT (**Figure S3H**). These data demonstrate that the magnitude of the Type I IFN response elicited by the SNP-7/8a vaccine can be augmented with ACT either T_scm_ or T_eff_ cells.

To extend these analyses, we performed a multiplexed ELISA for a large panel of cytokines and chemokines in serum and spleen following ACT with and without vaccination. There was a broad increase in GM-CSF, IL-12p40, TNFα, CXCL1, and IL-6 in mice that received T_scm_ cells and innate stimulation from either Ova:SNP-7/8a or E7:SNP-7/8a compared to mice receiving T_eff_ cells alone (**Figure 3H**). In addition, T cell derived cytokines IFN-γ and IL-2 were highest in mice that received T_scm_ cells + Ova:SNP-7/8a compared to OVA:SNP or E7:SNP-7/8a (**Figure 3H and 3I**). Three days following vaccination, there was also a significantly increased proportion of of CD8^+^ splenocytes producing IFN-γ in mice that received T_scm_ cells + Ova:SNP-7/8a compared to T_eff_ cells + Ova:SNP-7/8a (**Figure 3J**). These data suggest that the qualitative differences in the T cells transferred have demonstrable effects on both innate and adaptive cytokine production following vaccination with both antigen and innate stimulation.

### Transcriptomic analysis of the TME by scRNAseq

Therapeutic vaccination with various formulations can remodel the tumor microenvironment to mediate tumor control in a type I IFN-dependent manner^27^. However, TME remodeling following ACT has been underexplored, particularly in combination with vaccination. To investigate TME remodeling following ACT and vaccination, CD45^+^ immune cells were sorted from tumors and scRNAseq libraries were generated (**Figure 4A**). Unbiased UMAP clustering revealed stable populations of macrophages, monocytes, dendritic cells, basophils, neutrophils, T cells, NK cells, and B cells (**Figure 4B**). The relative prevalence of these populations varied significantly based on whether tumor received vaccination, while differences in transferred cell quality did not mediate major changes (**Figure S4A**). Notably, the monocyte/macrophage (MonoMac) clusters separated into six discrete subclusters: Mono 1, Mono 2, Mono 3, Mac 1, Mac 2, and Mac 3. The macrophage subclusters were distinguished by expression of *Ctsd*, *Hmox1*, *C5ar1*, and *Ecm1* (**Figure 4C**). The transcriptional profile of the Mac 1 subset was characterized by higher expression of *Nos2*, *Inhba*, *Il7r*, *Il1a*, *Pdpn*, and enriched for the Gene Ontology (GO) annotation terms Wound healing, Positive regulation of epithelial to mesenchymal transition, Tissue remodeling, and VEGF production (**Figures 4C and 4D**), characteristic of tumor resident macrophages. While the Mac 1 and Mac 2 populations displayed tumor resident and migratory macrophage transcriptional profiles, respectively, the Mac 3 phenotype was driven by response to type I IFN. Key differentially expressed genes in the Mac 3 population included the *Ifit3*, *Cd38*, and the serum amyloid gene *Saa3* (**Figure 4C**). By GO term enrichment, Mac 3 cells were identified by Response to Virus, Antigen presentation of exogenous peptide on MHC I, and cellular response to IFN-α (**Figure 4D**).

**Figure 4.**
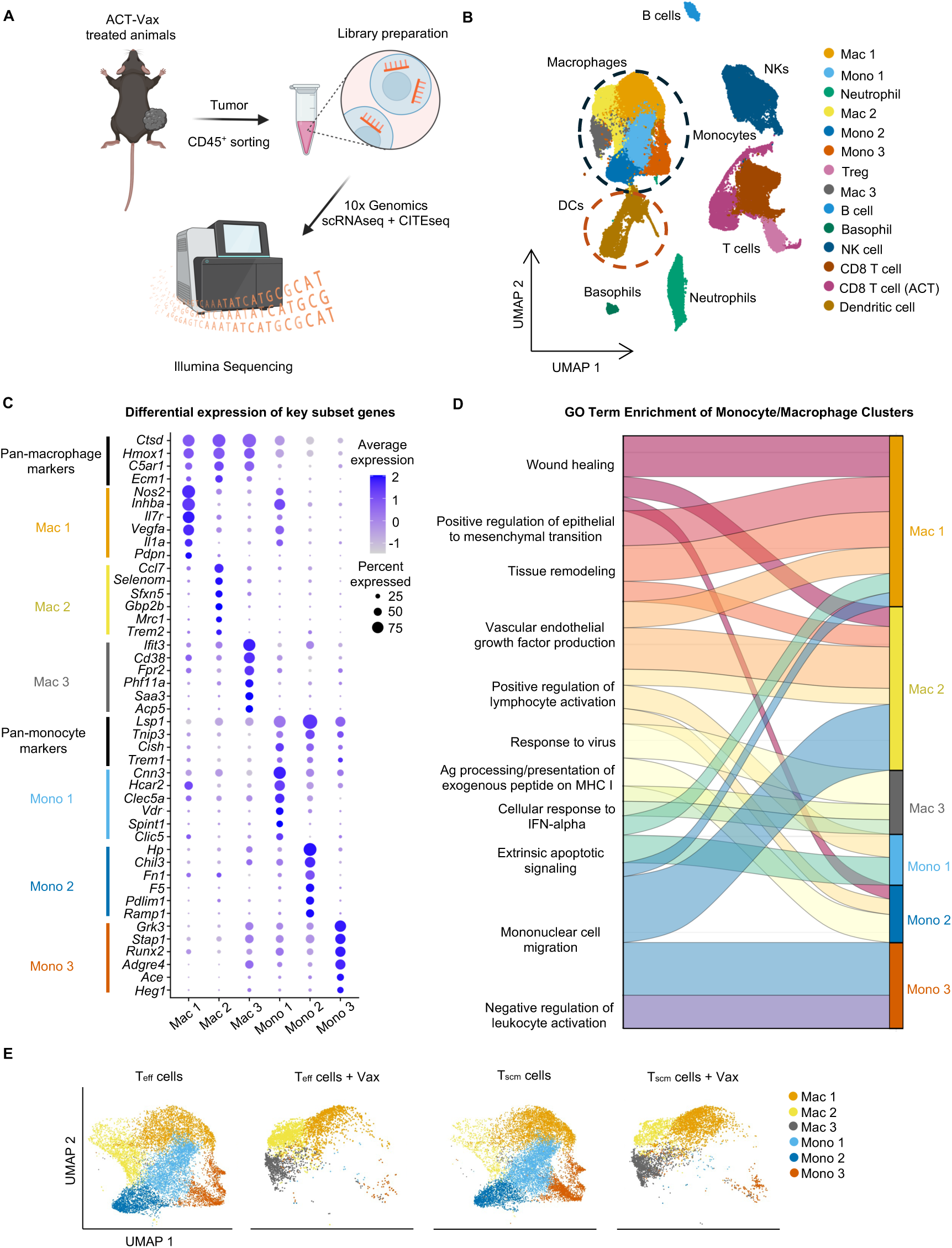
Transcriptomic analysis of the TME by scRNAseq. (A) Experimental schematic of harvesting and sorting CD45^+^ tumor cells from mice treated with ACT with or without vaccination, followed by generation of scRNAseq libraries. (B) UMAP detailing annotated clusters of immune cells present in the TME 24 hours following vaccination. The macrophage/monocyte cluster is circled in black and the dendritic cell cluster is circled in brown. (C) Dot plot highlighting differentially expressed genes used to define and annotate each cluster of monocytes and macrophages. Average and percent expression is indicated by size and color of dots. (D) Alluvial plot showing the GO term enrichments of each monocyte and macrophage cluster. Ribbon widths are proportional to the significance of GO term enrichment. (E) UMAP plots showing the monocyte and macrophage clusters present in indicated treatment group 24 hours following vaccination.

The immunoregulatory genes *Cnn3* and *Tnip3* are differentially expressed by all three monocyte subsets (**Figure 4C**). GO pathways enriched by the monocyte subsets include Positive regulation of lymphocyte activation and Extrinsic apoptotic signaling (Mono 1), Wound healing and Response to virus (Mono 2), and Mononuclear migration and negative regulation of leukocyte activation (Mono 3). The Mono 2 population contains a suppressive^27,50^ phenotype characterized by high differential expression of *Hp* and *Chil3*, a marker previously shown to have suppressive function in the TME. Notably, nearly all monocyte subsets were removed from the TMEs of mice receiving vaccination (**Figure 4E**). Conversely, the Mac 3 population is only found in vaccinated mice. The vaccine-induced increase in the Mac 3 population could be derived from one or multiple of the monocyte populations that are not detectable following vaccination or are recruited from blood monocytes^51^. To address these possibilities, two different computational analyses were performed. First, RNA velocity analysis via scVelo identified the Mono 2 population as a precursor population giving rise towards the other monocyte subsets (**Figure S4B**). Next, Slingshot trajectory analysis using the Mono 2 population as a starting population provided three trajectories in which Mono 2 cells differentiate to Mono 1 cells, which can then give rise to either Mono 3 cells, Mac 1 cells, or the vaccine-induced Mac 3 cells (**Figure S4C**). Thus, following ACT with either T_eff_ or T_scm_ cells, IV vaccination induced a pro-inflammatory Mac 3 cluster and eliminated all monocyte populations.

Dendritic cells can also have an active role in the tumor microenvironment by both stimulating T cells within the tumor and transporting tumor antigen to nearby lymph nodes to prime new anti-tumor responses via cross-presentation^52,53^. We subclustered the broad dendritic cell population which segregated into four distinct cell types: conventional dendritic cells (cDC1 and cDC2), plasmacytoid dendritic cells (pDC), and mature dendritic cells enriched in immunoregulatory molecules (mRegDC) (**Figures S4D and S4E**). There were minor differences in the composition of cDC1s and pDCs across treatment groups, with a modestly increased percentage of cDC1s in mice receiving T_scm_ cells as well as a small increase in the percentage of pDCs in unvaccinated mice (**Figure S4F**). More strikingly, vaccinated mice had significantly less cDC2s and a DC population largely composed of mRegDCs (**Figures S4F and S4G**). mRegDCs role in the TME has been associated with either immunosuppressive or anti-tumoral functions^54^. In clinical studies, mRegDCs have been associated with improved prognosis in patients with melanoma^55,56^.

Last, in terms of T cell analysis, both endogenous CD8^+^ T cells and transferred CD45.2^+^CD8^+^ T cells from the TME at this time point (24 hours after vaccination) were analyzed. There were no significant differences in the transcriptional phenotype or composition of T cells across the different treatment groups (**Figure 4B and S4A**). These data suggest that a longer timeframe is likely necessary to observe the differences in the T cell compartment shown by flow cytometry analysis three days after vaccination (**Figures 2C-2F**).

### Subcellular spatial transcriptomics of TME following ACT and vaccination

scRNAseq analysis of the TME 24 hours after vaccination showed similar immune compositions in animals that received vaccination following ACT with either T_eff_ or T_scm_ cells (**Figure S4A**). However, only mice that received T_scm_ cells + Vax had significant tumor protection (**Figure 2B**). Thus, to further analyze the potential *in situ* mechanisms of protection by T_scm_ cells + Vax, spatial transcriptomics on tumor sections was performed three days following vaccination, when there were significant differences in the magnitude and phenotype of T cells (**Figures 2C-2F**) and differences in tumor control between treatment groups emerged (**Figure 2B**).

Visium HD, which allows for unbiased whole transcriptome mRNA expression from subcellular 2×2μm resolution within a 6.5×6.5mm capture area^57^ was used for this analysis. To analyze the TME from spatial transcriptomics data, cells were segmented and 2×2μm barcodes that were contained within cell boundaries were assigned to respective cells (**Figure 5A and Methods**). Cell types annotated from UMAP clusters included tumor cells (stable tumor, progressive tumor, IFN-reactive tumor), macrophages (tumor resident, stimulatory, proliferative, proliferative stimulatory), monocytes, fibroblasts, epithelial cells, and T cells (**Figure S5A**). UMAP clusters were projected onto cell coordinates to form an image containing cell segments from the original H&E image labelled with cellular annotation (**Figure S5B**). Only a portion of the tumor section from the T_eff_ cells group was transcriptionally recovered, representing a small immune infiltrate on the periphery of the tumor that was largely comprised of tumor resident macrophages and proliferating tumor macrophages (**Figure S5B**).

**Figure 5.**
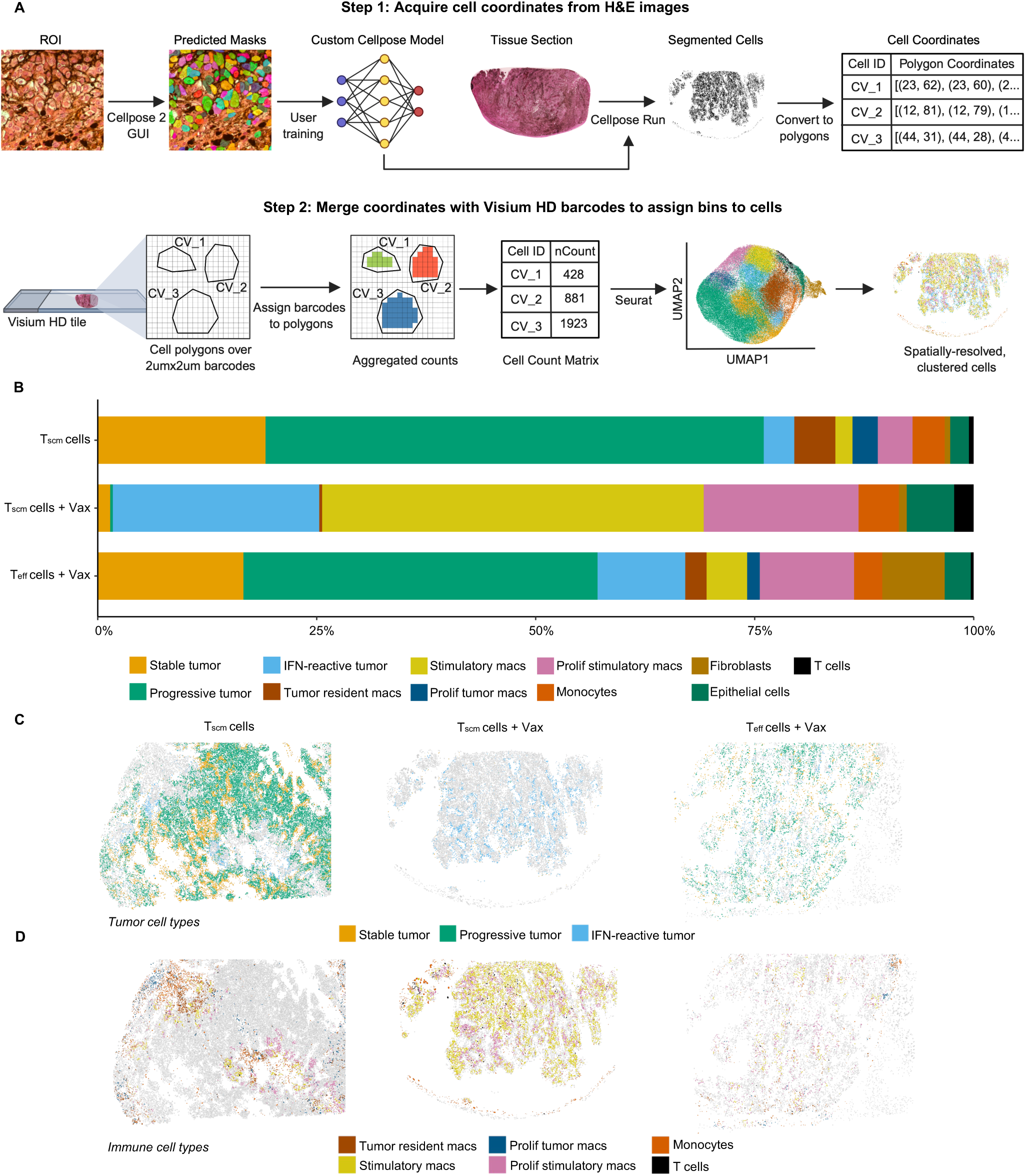
Subcellular spatial transcriptomics of TME following ACT and vaccination. (A) Schematic of computational pipeline to segment and derive spatial coordinates of cells used for spatial transcriptomics (top) and assign transcriptional barcodes to cluster and annotate spatially resolved cells (bottom). (B) Stacked bar graph showing the proportions of annotated cell types present within indicated treatment groups. (C) Spatial feature plots showing the presence and location of annotated tumor cell types within indicated treatment groups. (D) Spatial feature plots showing the presence and location of annotated immune cell types within indicated treatment groups.

Tumor cells were annotated based on the expression of tumor maintenance genes *Cdkn1a*, *Aldh1l2*, *Vegfa*, and the melanin-producing gene *Tyr* (**Figure S6A**). Notably, a population of “progressive” tumor cells were enriched for cell cycle genes like *Stmn1* and *Top2a*, indicative of high mitotic activity. IFN-reactive tumor cells had the highest expression of IFN response genes compared to other tumor clusters, including *Gbp2*, *Tap1*, and *Cxcl10*. In comparison with stable tumor and progressive tumor, IFN-reactive tumor was enriched in Hallmark pathways that included IFN-γ response, IFN-α response, TNFα signaling via NFκB, and Apoptosis (**Figure S6B**). Apoptotic genes differentially expressed in IFN-reactive tumor included genes involved in both intrinsic and extrinsic apoptosis, including *Tap1*, *Jun*, *Cflar*, *Casp7*, and *Bid* which are associated with T cell mediated killing^58^ (**Figure S6C**). Of note, there were major differences found in the prevalence of different of tumor cell phenotypes within the TME based on treatment group. The relative proportion of stable tumor cells and progressive tumor cells were increased in tumors that received T_scm_ cells alone or T_eff_ cells with or without vaccination (**Figure 5B and S6D**). Strikingly, tumors from animals treated with T_scm_ cells + Vax were largely composed of IFN-reactive tumor cells (**Figures 5B, S6D**). Contiguous sheets of tumor cells were found throughout tumor specimens, however, the spatial distribution of immune cells varied by treatment group (**Figures 5C and 5D**).

Heterogenous clusters of macrophage cells were also observed in the TME and annotated based on expression of *H2-Ab1*, *Axl*, and *Cd74* (**Figure S6A**). Two pro-inflammatory subsets of macrophages (stimulatory macrophages and proliferating stimulatory macrophages) were enriched for *Stat2*, *Ifit2*, *Irf8*, *Oasl2*, and *Ciita* (**Figure S6A**). Compositionally, tumors treated with T_scm_ cells alone contained a smaller proportion of stimulatory macrophages, while the immune compartment of T_scm_ cells + Vax was overwhelmingly composed of stimulatory macrophages (**Figure S6E**). These data highlight treatment-dependent differences in both the composition and relative frequency of cell types that exist in the TME following ACT, depending on the quality of transferred cells and vaccination status.

### Effect of intravenous vaccination on an immune-active TME

To examine the spatial relationship between immune cells and tumor cells across treatment groups, spatial graphs were divided into rectangles containing ∼25 cells per area. Areas containing at least one tumor cell and at least one immune cell were highlighted in red (**Figure S7A**). To visualize how tumor-immune interactions were related to regions of the TME with the highest amounts of tumor cells, heatmaps were projected describing the proportion of tumor cells (**Figure S7B**) and immune cells (**Figure S7C**) in each rectangle. The merging of the tumor density and tumor-immune interaction graphs revealed that following ACT with T_scm_ cells + Vax or T_eff_ cells + Vax, immune cells penetrated throughout the parenchyma and were juxtaposed to tumor cells, indicative of an “immune active” TME (**Figure 6A**). In contrast, ACT with T_scm_ cells alone showed an “immune excluded” TME, where the tumor-immune barrier is highly demarcated, separating areas containing the highest density of tumor cells from surrounding immune cells (**Figure 6A**). The difference in total tumor counts versus only those contained within red tiles reveals additional data on the tumor-immune interactions. With T_scm_ cells alone, there is a striking reduction in stable and progressive tumor cells when viewing only tumor cells spatially associated immune cells; however, 91% of IFN-reactive tumor cells are contained within the red tiles, suggesting a correlation between immune cells and the IFN-reactive tumor phenotype (**Figure S7D**). Importantly, due to immune infiltration following intravenous vaccination, nearly all tumor cells are contained within red tiles in both the T_eff_ cells + Vax and T_scm_ cells + Vax treatments, despite significant differences in the proportion of tumor phenotypes in those treatment groups (**Figures S6D and S7D**). The composition of immune cells contained within red tiles also varied by treatment group, with the TME treated with T_scm_ cells alone containing a smaller proportion of stimulatory and proliferative stimulatory macs than those that were vaccinated (**Figure S7E**). These data demonstrate an important effect to drive immune infiltration in an otherwise immune-excluded tumor by combining ACT with intravenous vaccination.

**Figure 6.**
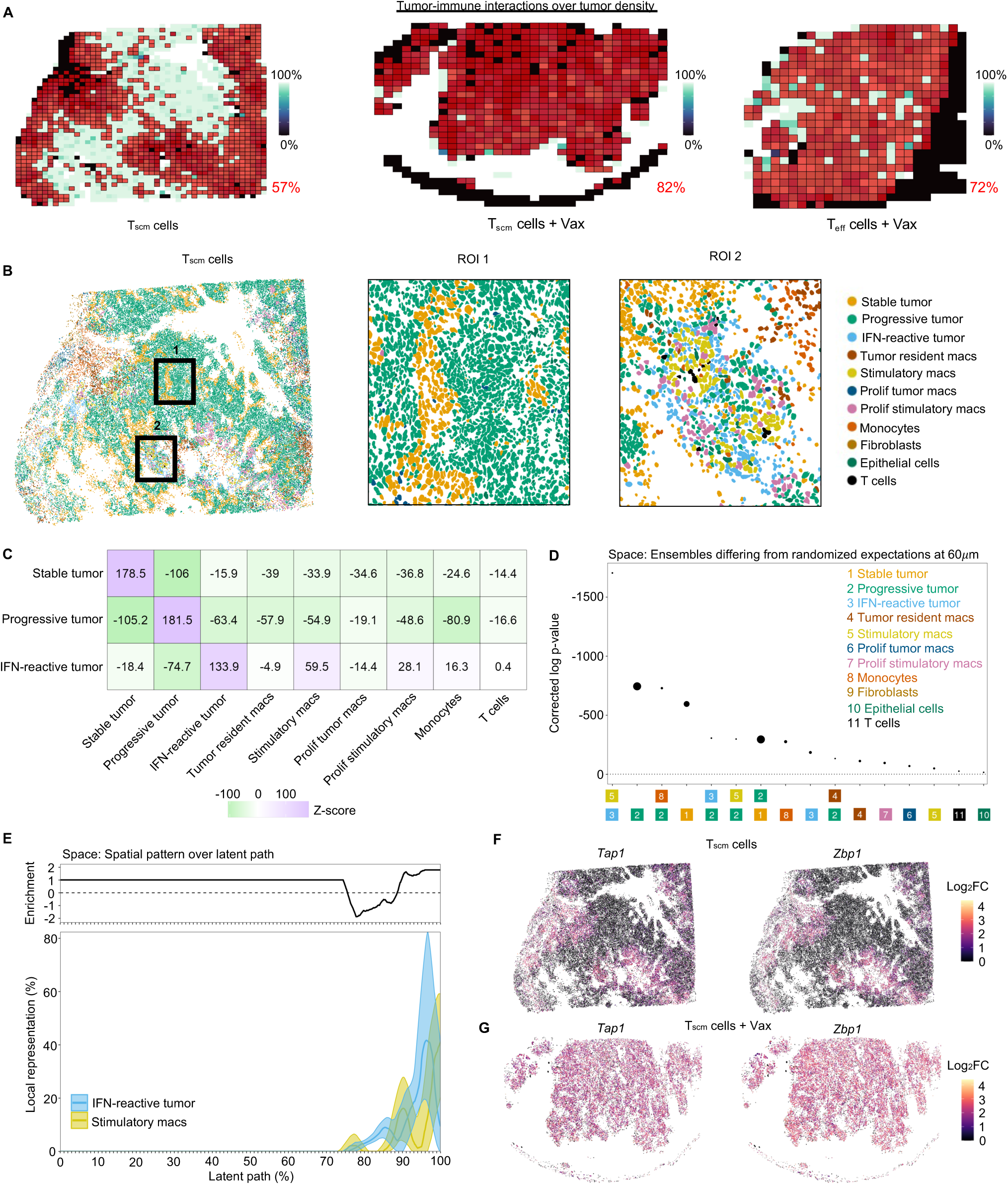
Tumor-immune cell interactions by spatial transcriptional analysis. (A) Spatial feature plots of indicated treatment groups showing the presence of tumor-immune interactions in local tiled neighborhoods (red) overlaying scaled heatmap of tumor cell density within each tiled neighborhood. (B) Spatial feature plot of segmented cells in TME treated with Tscm cells alone. Black boxes indicate areas of tumor parenchyma composed of tumor cells (ROI 1) or tumor periphery composed of mixture of tumor cells and immune cells (ROI 2). (C) Heatmap showing the Z-scores of neighborhood enrichment analysis based on proximity, highlighting high (purple) or low (green) enrichment between neighbors. (D) Space plot showing significant patterns of cellular ensembles within the TME ranked by corrected log p-value. Point size is proportional to cisMI magnitude. (E) Covariation plot detailing the smoothed mean and 95% confidence interval of indicated cell local representation across latent path with running enrichment score. (F) Spatial feature plots of TME treated with Tscm cells highlighting the Log2FC expression of *Tap1* and *Zbp1* within each cell. (G) Spatial feature plots of TME treated with Tscm cells + Vax highlighting the Log2FC expression of *Tap1* and *Zbp1* within each cell.

### Tumor-immune cell interactions by spatial transcriptional analysis

The finding that the tumor transcriptional phenotype is heterogenous and affected by spatial proximity to immune cells led to further analysis to assess which spatial relationships may be important for the differences observed in tumor composition across treatment groups (**Figure S6D**). By magnifying regions of interest (ROI) within the TME of the animals that received T_scm_ cells alone showed few (ROI 1) or many (ROI 2) tumor-immune interaction tiles. There were progressive and stable tumor cells in ROI 1 (**Figure 6B**). In contrast, ROI 2 has a larger proportion of IFN-reactive tumor cells, surrounded by stimulatory macs, proliferative stimulatory macs, and T cells (**Figure 6B**). Following ACT with either T_eff_ cells or T_scm_ cells and vaccination, the TME was more spatially homogenous (**Figure S8A and S8B**). To statistically determine the enrichment of tumor cells with other cell types in the TME treated with T_scm_ cells, we applied a k-nearest-neighbor (kNN) algorithm followed by neighborhood enrichment to calculate z-scores (**Figure 6C**). While each tumor cell phenotype strongly associated with other cells of the same phenotype, only IFN-reactive tumor cells were positively associated with immune cells including stimulatory macs, proliferative stimulatory macs, monocytes, and T cells. Notably, tumor resident macs and proliferative tumor macs did not spatially associate with IFN-reactive tumor cells (**Figure 6C**). To further investigate the spatial patterns that exist within the TME, spatial patterning analysis of cellular ensembles^59^ (SPACE) was used. A census of 60μm neighborhoods within the image revealed several significant ensembles, including stimulatory macrophages with IFN-reactive tumor cells and monocytes with progressive tumor cells (**Figure 6D**). Notably, IFN-reactive tumor did not form a significant ensemble with any other immune cell. By observing the cell types across a “latent path” representing the highest density spatial neighborhoods, it is notable that in every instance where IFN-reactive tumor is present, it is co-localized with stimulatory macrophages (**Figure 6E**). Moreover, by graphing the latent path of IFN-reactive tumor with all immune cells, only stimulatory macrophages are co-enriched with IFN-reactive tumor, each peaking ∼95% through latent path (**Figure S8C**). Of note, stimulatory macrophages also formed an ensemble with a small proportion of the progressive tumor cells (**Figure 6D**). Plotting their co-representation along the latent path revealed a pattern that in the neighborhoods with the least proportion of progressive tumor cells (around the periphery), there is a co-enrichment with stimulatory macrophages that also spatially reside in this area of the tissue (**Figure S8D**). Due to the significant spatial relationship between stimulatory macrophages and IFN-reactive tumor cells, it is possible that IFN-reactive tumor cells are being converted to this phenotype from progressive tumor cells. This is also suggested by the significant ensemble formed between IFN-reactive tumor cells and progressive tumor cells (**Figure 6D**), where the neighborhoods with the highest proportion of IFN-reactive tumor cells are co-enriched with progressive tumor cells as well as stimulatory macrophages.

Last, to determine transcriptional changes associated with spatial patterns, spatially differentially expressed genes, including the antigen presentation gene *Tap1* and *Zbp1*, which has been shown to promote tumor immunogenicity and necroptosis^60,61^. *Tap1* and *Zbp1* were spatially restricted to areas containing stimulatory macrophages and IFN-reactive tumor in the TME treated with T_scm_ cells alone (**Figure 6F).** In contrast, these transcripts were ubiquitously expressed across the TME treated with T_scm_ cells + Vax (**Figure 6G**). Within areas containing progressive and stable tumor, the cell cycle genes *Mki67* and *Tacc3*, which has been associated with tumor aggression and poor prognosis, are differentially expressed (**Figure S8E**). These data detail a highly significant spatial association between stimulatory macrophages and IFN-reactive tumor cells and suggest a mechanism in which proximal interaction between stimulatory macrophages and progressive tumor cells can drive transcriptomic remodeling towards an immunogenic tumor phenotype.

### Spatial transcriptomic differences in TME remodeling following ACT and vaccination

By scRNAseq analysis 24 hours following vaccination, the immune composition of the TME was highly similar between T_eff_ cells + Vax and T_scm_ cells + Vax (**Figure S4A**), suggesting that IV vaccination is sufficient to remodel the TME following ACT with T_eff_ or T_scm_ cells. However, the spatial transcriptomic analysis of samples taken three days following vaccination showed significant differences between the tumor and immune compositions by ACT with T_eff_ and T_scm_ cells (**Figures 5B, S6D, and S6E**), demonstrating a major effect that the quality of transferred cells can have on the structure of the TME. Thus, to further examine how T cell quality impacts TME dynamics, we performed subsequent analyses comparing T_eff_ cells + Vax and T_scm_ cells + Vax. As stimulatory macs were spatially enriched near IFN-reactive tumor in the T_scm_ cells TME (**Figures 6C-6E**), the density of stimulatory macs across the TMEs treated with T_scm_ or T_eff_ cells and vaccination were analyzed (**Figure 7A**). Tumor treated with T_scm_ cells + Vax had a significantly higher prevalence of stimulatory macs per bin (**Figure 7B**). This is consistent with the higher proportion of IFN-reactive tumor cells following treatment with T_scm_ cells + Vax compared to T_eff_ cells + Vax (**Figure 5B and S6D**). As proliferative stimulatory macs are more represented in the T_eff_ cells + Vax TME and have a similar phenotype to stimulatory macs and a visible (albeit non-significant) spatial association with IFN-reactive tumor cells (**Figure S8C**), we also examined the prevalence of both phenotypes. Even when controlling for the addition of proliferative stimulatory macs, the T_scm_ cells + Vax TME still contained a significantly greater density of immunostimulatory macrophages (**Figure S9A and S9B**). There was also a higher density of T cells following T_scm_ cells + Vax (**Figure S9C and S9D**). To assess how differences in cellular composition across the TMEs affect global transcriptomic signatures, we performed a differential gene expression analysis followed by enrichment for Hallmark gene sets. We found that the TME treated with T_scm_ cells + Vax was highly enriched for pro-inflammatory immune pathways while treatment with T_eff_ cells + Vax yielded enrichment for tumor progression pathways such as mitotic spindle and angiogenesis (**Figure 7C**). Indeed, 100 DEGs (*P* < 0.05 and log_2_FC > 1, Wilcoxon test) were identified with genes related to wound healing upregulated in T_eff_ cells + Vax and genes related to immune receptors, cytokine signaling, interferon response, metabolic signaling, and antigen presentation upregulated in T_scm_ cells + Vax (**Figure S10A**). GSEA showed high enrichment for IFN-γ and IFN-α response pathways following T_scm_ cells + Vax treatment (**Figure 7D**) and high enrichment for epithelial-to-mesenchymal transition and myogenesis following T_eff_ cells + Vax treatment (**Figure 7E**), pathways associated with tumor metastasis^62^. These data demonstrate a major difference in TME remodeling that can be achieved by using stem-like CD8^+^ T cells for ACT.

**Figure 7.**
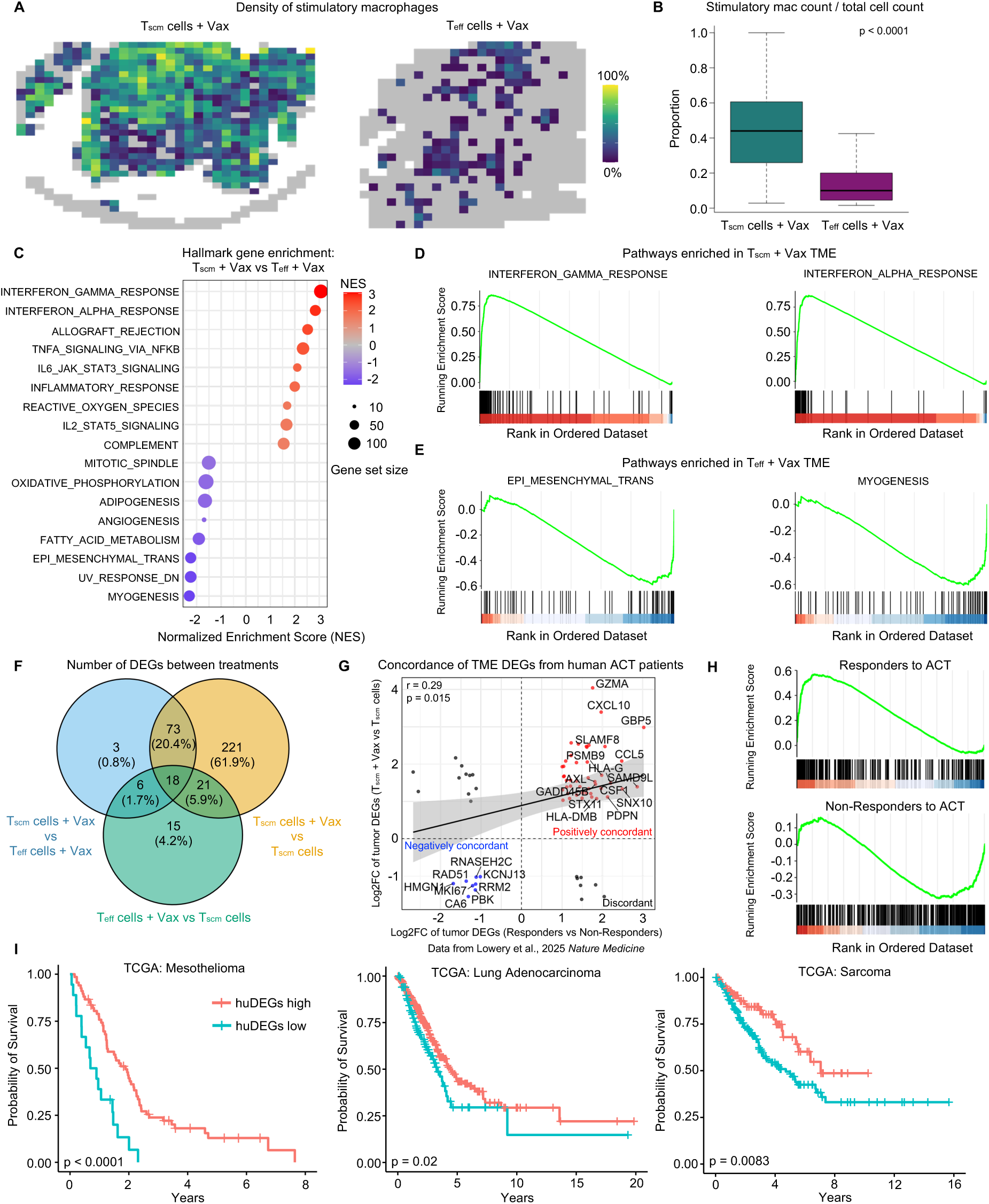
Spatial transcriptomic differences in TME remodeling following ACT and vaccination. (A) Spatial feature plots of indicated treatment groups displaying a scaled heatmap of stimulatory macrophage cell density within each tiled neighborhood. (B) Bar graph quantitating the proportion of stimulatory macrophages per tiled neighborhood in tiles containing at least one stimulatory macrophage in indicated treatment groups. Statistics accessed by a two-sided Wilcoxon test. (C) Dot plot indicating Hallmark gene pathways that are positively (red) or negatively (blue) enriched by comparing TMEs treated with Tscm cells + Vax and Teff cells + Vax. (D) Enrichment plots showing running enrichment scores in the Hallmark Interferon Gamma Response (left) or Interferon Alpha Response (right) gene sets across ranked DEGs comparing TMEs treated with Tscm cells + Vax and Teff cells + Vax. (E) Enrichment plots showing running enrichment scores in the Hallmark Epithelial to Mesenchymal Transition (left) or Myogenesis (right) gene sets across ranked DEGs comparing TMEs treated with Tscm cells + Vax and Teff cells + Vax. (F) Venn diagram showing the number of overlapping and non-overlapping DEGs calculated by TME comparisons of Tscm cells + Vax vs Teff cells +.Vax (blue), Tscm cells + Vax vs Tscm cells (yellow), and Teff cells + Vax vs Tscm cells (green). (G) Correlation plot graphing genes matched from the TMEs of human responders to ACT and humanized genes from the TME following Tscm cells + Vax treatment. Genes plotted by log2FC expression compared to non-responders. Genes were labelled as positively concordant (red) if they were upregulated in both studies, negatively concordant (blue) if they were downregulated in both studies, or discordant (black) if they had different regulations between studies. Statistics accessed by Pearson correlation test. (H) Enrichment plots showing running enrichment scores in genes upregulated in responders to ACT (top) or non-responders to ACT (bottom) across ranked, humanized DEGs comparing TMEs treated with Tscm cells + Vax and Tscm cells. (I) Kaplan-Meier survival curves from TCGA patient data of Mesothelioma (left), Lung Adenocarcinoma (middle), and Sarcoma (right). Patient cohorts were stratified as high (red) or low (blue) based on huDEGs signature score. Statistics accessed by log-rank test.

### Transcriptomic TME signatures in mice and humans with improved outcomes

In advanced tumors, where the TME may negatively impact the effectiveness of T cell-based therapies, identifying transcriptional biomarkers to predict therapeutic outcomes or guide therapies to modulate the TME may be critical. To determine the transcriptional biomarkers that differentiated the TMEs in this murine model, DEG analysis was performed by pseudobulking TME spatial transcriptomic data and comparing global RNA expression between each treatment group, thereby revealing which genes were altered by ACT with T_eff_ cells, T_scm_ cells, or vaccination status (**Methods**). The largest number of DEGs were found by comparing the TMEs of mice treated with T_scm_ cells versus T_scm_ cells + Vax. This analysis showed 333 DEGs due to the effect of the vaccine on mice treated with T_scm_ cells (**Figure 7F and Supplementary Table 1)**. 100 DEGs defined differences between mice treated with T_scm_ cells + Vax and T_eff_ cells + Vax (**Figures 7A and S10A**), which described the effect of transferred cell quality on the TME. Lastly, as mice treated with T_scm_ cells or T_eff_ cells + Vax had similar tumor control and tumor cell composition at this timepoint (**Figures 2B and S6D**), DEG analysis was performed between these groups to investigate gene programs that were insufficient in controlling tumor growth. Remarkably, despite receiving different quality T cells for ACT and differing in vaccination status, only 60 genes differentiated the TMEs treated with T_scm_ cells and T_eff_ cells + Vax (**Figure 7F**). 39 of the 60 genes were also differentially expressed between T_scm_ cells and T_scm_ cells + Vax, indicative of transcripts that were upregulated by vaccination that were not sufficient to drive therapeutic effect (**Supplementary Table 1**). These data suggest that T_scm_ cells are capable of a significantly greater modulatory effect on the TME following vaccination compared with T_eff_ cells.

To determine if the transcriptomic tumor signature following ACT with T_scm_ cells + Vax was predictive of clinical transcriptional responses from humans with cancer, we compared DEGs from TME treated with T_scm_ cells + Vax with bulk RNAseq tumor signature data of responders versus non-responders in a recent clinical trial of TIL therapy in gastrointestinal cancer^63^ (**Figure 7G**). After humanizing the T_scm_ cells + Vax tumor DEGs and plotting their log2FC expression versus the log2FC expression of matched genes differentially expressed in the responder patient tumors, 53/71 (75%) genes were positively or negatively concordant while 18/71 (25%) genes were discordant (r = 0.29*, P* = 0.015). By GSEA of ranked humanized T_scm_ cells + Vax tumor DEGs (**Figure 7H**), responders to ACT were positively enriched (NES = 2.21, *P* < 0.0001) while non-responders to ACT were negatively enriched (NES = -1.56, *P* < 0.0001).

To determine whether the T_scm_ cells + Vax tumor signature was predictive of survival across several different human cancers, patients in the cancer genome atlas (TCGA) were scored for high or low enrichment of the humanized DEGs (huDEGs) defined here (**Supplementary Table 2**). Across all tumors represented in the TCGA, patients enriched for the huDEG signature had improved survival (**Figure S10B**). Individual tumors with significantly improved survival included mesothelioma, lung adenocarcinoma, and sarcoma (**Figure 7I**). Some less common tumors had improved survival when enriching for the huDEGs (**Figure S10C-S10E**), including adenoid cystic carcinoma, kidney chromophobe, and papillary renal cell carcinoma. These data show that the transcriptomic signature associated with the TME of T_scm_ cells + Vax treated tumors in this murine model may also be predictive of responsiveness in a human cohort of ACT patients and associated with improved survival in a pan-tumor atlas of human cancer patients.

## Discussion

ACT is a promising approach for treating advanced tumors, with clinical outcomes determined by several critical steps. Here, in a murine tumor model, we developed a comprehensive approach to improve ACT therapy at each step and then assessed the *in situ* transcriptional dynamics of immune cells within the TME by spatial transcriptomics. The primary findings of this study are the following: 1) Naïve murine CD8^+^ T cells activated in the presence of CAL-101 retain a stem-like phenotype, expand *in vivo* following intravenous vaccination, and mediate tumor control 2) Intravenous vaccination elicits systemic innate immunity that can modulate the TME and augments systemic cytokine and chemokine production following ACT with T_scm_ cells 3) Spatial transcriptomics reveals that IV vaccination converts an immune-excluded tumor into an immune-infiltrated tumor, and identifies a spatial relationship between stimulatory macrophages and IFN-reactive tumor cells 4) Transcriptomic biomarkers from the TME of mice that controlled tumor growth was concordant with TME analysis of humans that responded to ACT or that had improved survival outcomes to a broad range of tumors.

The first step toward optimizing ACT was focused on generating stem-like CD8^+^ T cells, a population based on their enhanced durability and improved effector functions against tumors^9^. Consistent with prior studies^15^, the PI3k inhibitor CAL-101 promoted the generation of stem-like CD8^+^ T cells during *in vitro* culture. These T_scm_ cells expressed the canonical stem-like markers *Tcf7*, *Sell*, and *Slamf6* (**Figure 1H**). There was also a unique cluster of T_scm_ cells containing the highest expression of *Tcf7* and *Sell* that also differentially expressed *Pik3ip1*, a negative regulator of PI3k, compared to other stem-like clusters, implying a mechanism by which cells receiving the greatest inhibition of PI3k retain the largest stem-like characteristics (**Figure 1K**). Of note, despite being associated with PI3k inhibition^64^ the transcription factor *Bach2*, which has been shown to be a major regulator of T cell differentiation and associated with T_scm_ cells, was not differentially expressed in cells treated with CAL-101. These data underscore the diverse approaches to sustaining stem-like CD8^+^ T cells by differentially modulating T cell signaling pathways. The ability of CAL-101 to target PI3k signaling within T cells and prevent differentiation towards effector cells suggests it could also be used in maintaining stem-like T cell quality across different ACT approaches, such as TCR, CAR, and TIL therapy. Indeed, the finding that TIL therapy with a higher frequency of CD39^-^CD69^-^ T cells, markers associated with stem-like quality, were correlated with improved clinical response^65^ provide additional evidence for this optimization approach.

Following ACT, a standard approach for expanding T cells *in vivo* is to condition the recipient through radiation or chemotherapy to induce lymphodepletion allowing for homeostatic expansion^66^. Additionally, IL-2 is also used following ACT, most commonly with TILs, for T cell expansion^1,67^. However, these interventions can lead to clinical toxicity^68,69^, and high dose IL-2 may also expand regulatory T cells^70^. Here, we used vaccination to expand transferred T cells without any pre-conditioning step. Moreover, ACT was done with a relatively small magnitude of CD8^+^ T cells (10^5^) compared to other studies using ACT in a similar tumor model^71,72^. The ability to use stem-like cells for ACT may allow for lowering the magnitude needed to achieve tumor control. Vaccination following ACT resulted in a higher magnitude of transferred T_scm_ cells in the tdLN and spleen compared to T_eff_ cells, as well as significantly improved tumor control (**Figures 2B and 2C**). These data are consistent with the improved function of stem-like CD8^+^ T cells and may help inform clinical studies aiming to improve ACT with vaccination^73^. Vaccination to expand the cognate antigen expressed by the transferred T cells may also lead to expansion of endogenous tumor-specific T cells. The finding that only T_scm_ cells were able to regress a mixed tumor with only 50% of cells expressing target antigen following vaccination is consistent with antigen spreading^48,74^.

A key finding of this study was to demonstrate the differences between ACT using T_scm_ and T_eff_ cells for tumor control. In our analysis of serum cytokines in mice treated with T_scm_ cells or T_eff_ cells and vaccination, there were notable differences in the amount of both innate and adaptive cytokines. Higher serum concentrations of IFN-γ, IL-2, IL-12p40, TNFα, and CXCL1 were observed following ACT with T_scm_ cells + Vax compared to T_eff_ cells + Vax (**Figure 3H and 3I**). Furthermore, there were significantly more IFN-γ producing CD8^+^ T cells in the spleens of mice treated with T_scm_ cells + Vax than T_eff_ cells + Vax three days following vaccination (**Figure 3J**). The enhanced production of IFN-γ and TNFα, canonical mediators of tumor killing that have been shown to mediate improved clinical responses following ACT^75^, confirm functional differences *in vivo* between ACT with T_eff_ and T_scm_ cells followed by vaccination.

Intravenous vaccination with the SNP-7/8a vaccine can also have direct effects on the immune composition of the TME, by reducing the frequency of suppressive myeloid cells in a type I IFN-dependent manner^27^. Targeting suppressive monocyte and macrophage populations in the TME can enhance the function of intratumoral T cells and improve therapy with therapeutic cancer vaccines and CPI therapy^76–78^. Here, a notable finding was that systemic type I IFN induced by IV SNP-7/8a is enhanced when vaccination is combined with ACT compared to vaccination alone (**Figures 3E-3F and S3F-S3G**). One potential mechanism for the increased innate immune activation is T cell derived IFN-γ production, leading to enhanced CXCL9/CXCL10 signaling and recruitment of IFN-I producing myeloid cells^79^. IV vaccination following ACT with T_scm_ or T_eff_ cells also had a profound effect on the transcriptional and phenotypic profile of the TME with reduction of multiple monocyte populations and induction of a stimulatory Mac 3 population when assessed 24 hours later (**Figure 4E**). In this analysis of the TME, there were no differences between T_scm_ and T_eff_ cells following vaccination (**Figure S4A**). However, T_eff_ cells + Vax showed significantly less tumor control than T_scm_ cells + Vax (**Figure 2B**). Thus, early remodeling of the TME induced by the vaccination was not sufficient to optimize tumor protection. These data suggest that the T_scm_ cells are inducing additional changes in the TME over time to mediate tumor control.

To assess the *in situ* TME effects of ACT with T_scm_ versus T_eff_ cells on tumor and immune cell populations, we performed spatial transcriptomics three days after vaccination. Consistent with reports of B16-Ova as an immune-excluded tumor^80^, mice treated with T_scm_ cells alone had distinct separation between the tumor parenchyma and surrounding immune cells (**Figure 6A**). In striking contrast, TMEs treated with T_scm_ or T_eff_ cells and vaccination were immune-active, with infiltration of macrophages, monocytes, and T cells throughout the tumor (**Figures 6A, S7A-S7E**). Moreover, treatment with T_scm_ cells + Vax resulted in significantly greater densities of stimulatory macrophages and T cells throughout the TME compared to treatment with T_eff_ cells + Vax (**Figures 7A, S9A-S9D**). Lymphocyte- and myeloid-recruiting chemokines *Cxcl10* and *Ccl5* were overexpressed in the TME following T_scm_ cells + Vax compared to T_eff_ cells + Vax (**Figure S10A**), consistent with immune infiltration^81^.

Tumor cell heterogeneity and immunogenicity are critical prognostic factors in cancer therapy but have been underexplored in ACT models^82–84^. A notable finding from this study is the heterogeneity of tumor cells following the different ACT therapies, including an IFN-reactive tumor type. Hallmark gene enrichment of the TME following T_scm_ cells + Vax (comprised of >93% IFN-reactive tumor) showed enrichment for IFN-γ response, IFN-α response, allograft rejection, IL-6 signaling, and TNFα signaling (**Figure 7C**), the same top pathways enriched in the TMEs from the TCGA with the highest antigen processing and presenting machinery score (APS) used to measure tumor immunogenicity^83^. Thus, the transcriptomic signature of IFN-reactive tumor is consistent with enhanced immunogenicity as well as increased apoptotic signaling (**Figure S6C**). As spatial patterns of cells within a tumor can reveal important biological relationships^85,86^, we utilized an information-theoretic algorithm^59^ SPACE to discern where IFN-reactive tumor cells reside within the TME. Despite the TME being comprised largely of progressive and stable tumor cells following treatment with T_scm_ cells alone, SPACE identified a highly significant association between stimulatory macrophages and IFN-reactive tumor cells (**Figures 6D and 6E**). Importantly, IFN-reactive tumor cells were only observed when they were spatially associated with stimulatory macrophages (**Figure 6B**), suggesting a potential mechanism by which tumor cells can be converted towards an IFN-reactive state. Moreover, the TME following treatment with T_scm_ cells + Vax was comprised of the highest proportion of both stimulatory macrophages and IFN-reactive tumor amongst all treatment groups (**Figures S6D and S6E**). These data highlight a potentially powerful strategy to enhance tumor cell susceptibility to cell death.

The final analysis was to determine whether the transcriptional signatures in the B16 melanoma tumor model could be associated with transcriptional TME signatures in human patients that responded to ACT therapy. In comparing TME transcriptomic signatures following T_scm_ cells + Vax treatment, there was significant concordance with a recent clinical study comparing the TMEs of responders and non-responders to TIL therapy^63^ (**Figure 7F**). Genes that were associated with protection in both studies included anti-tumoral programs like Cytotoxicity (GZMA, STX11), IFN response (GBP5, IRF7, IFIT3), and chemokine signaling (CXCL10, CCL5) (**Supplementary Table 2**). The ability to define transcriptomic biomarkers that may predict outcome to ACT or other immune treatments may be useful for identifying patients likely to benefit from ACT or require additional intervention before or during ACT to maximize efficacy in the TME.

In conclusion, ACT is a powerful clinical approach that can regress established and metastatic solid tumors. Here we show that ACT with T_scm_ cells combined with IV vaccination improves T cell function and cytokine production *in vivo* and synergistically remodels the TME to enhance immune infiltration and tumor responsiveness to immunotherapy.

## Limitations

There are several limitations in this study. First, naïve transgenic-TCR mouse CD8^+^ T cells were used, which is standard for ACT models of tumor in mice. Application of the cell culture methods described here to maintain stemness from naïve cells may show differences in TCR T cells, CAR-T cells, and TILs that begin in a more differentiated state. Ongoing studies using mouse and human TILs are underway to extend the potential of using CAL-101 when starting with more differentiated T cells. The melanoma and colon tumors used in the models shown here may not represent the heterogeneity of the TME and immunoediting observed across human tumors. However, as the aims of this study were to interrogate experimental immunotherapy regimens and provide understanding of how ACT and vaccines are mediating protection under highly controlled conditions, the data generated may inform therapeutic approaches in humans. Last, we have previously shown that therapeutic neo-antigen vaccination with SNP-7/8a provides protection against tumors in a Type I dependent manner. While we showed that ACT with T_scm_ cells followed by vaccination with SNP-7/8a induced Type 1 and Type 2 cytokines *in vivo*, we did not formally show that they are required for protection. Ongoing studies are underway to provide confirmation of their role for tumor control in our studies and other models.

In terms of the spatial analysis, leakage of transcripts from cells to neighboring spots is a well-known technical limitation within spatial transcriptomic analysis. In our study, transcript leakage was noted by observing background transcript counts in areas not containing cells or not containing tissue (**Figures S10F and S10G**). This results in the contamination of cells with neighboring cell mRNA transcripts. Similar to analysis of ambient RNA in droplet-based sequencing methods, the most highly expressed transcripts of the most abundant cell populations contaminate the least abundant, neighboring cell populations^87,88^. For example, highly expressed *H2-Ab1* transcripts from abundant macrophages populations leaked into less abundant cell types: monocytes, fibroblasts, epithelial cells, and T cells (**Figure S6A**). However, leakage from T cell-specific genes (*Gzmb*, *Ifng*, *Cd8a*) were not observed as noticeable artifacts in macrophage clusters, due to their high transcript counts. Nevertheless, diverse cell types can still be annotated and spatially analyzed despite the presence leaked transcripts. Ongoing development of tools capable of regional transcript background subtraction will mitigate this technical issue.

## Methods

### Mice

Wild-type female 6-8 week old mice of strains C57BL/6J, B6.SJL-Ptprca Pepcb/BoyJ, C57BL/6-Tg(TcraTcrb)1100Mjb/J, and B6.Cg-Thy1a/Cy Tg(TcraTcrb)8Rest/J mice were purchased from The Jackson Laboratory and housed in specific-pathogen-free conditions. Mice were allowed to adjust to facility conditions for one week before experimental use. All experiments were performed in the vivarium of the Vaccine Research Center at the National Institutes of Health. All experiments were approved by the Institutional Animal Care and Use Committee at the NIH. All experiments were conducted under the ethical guidelines set by the Institutional Animal Care and Use Committee and animals were humanely euthanized at predefined endpoints.

### Tumor cell lines and implantation

MC38 cell line was a gift from L. Delamarre (Genentech). B16-gp100-KVP and MC38-gp100-KVP cell lines were a gift from Zhiya Yu (NCI). B16-OVA was a gift from H. Levitsky (Juno Therapeutics). Tumor cell lines were grown under standard conditions as previously described^45^ and were grown from frozen stocks for four days, and were washed, resuspended in sterile PBS, and filtered prior to implantation. 1×10^5^ B16-Ova or 3.5×10^5^ B16-gp100-KVP were implanted subcutaneously into shaven mice flanks. For experiments studying the tumor ratio of MC38-gp100-KVP:MC38, 1×10^5^ MC38-gp100-KVP tumor cells were inoculated per mouse for the 100:0 ratio, while MC38-gp100-KVP and WT MC38 tumor cells were mixed in a 50:50 mixture prior to inoculation before 1×10^5^ mixed cells were inoculated subcutaneously. Tumors were measured every 3-4 days using digital calipers and were euthanized when tumor size exceeded 1000mm^3^.

### *in vitro* T cell culture

WT OT-1 or p-mel mice spleens were harvested and passed through a 40-micron filter. Red blood cells were removed from splenocytes by ACK lysis and washed with PBS. Cells were resuspended in T cell culture medium (RPMI 1640 Medium containing 10% FBS, 1% penicillin/streptomycin/glutamine, 1% non-essential amino acids, 0.1% β-mercaptoethanol, 25mM HEPES, 1mM sodium pyruvate) and supplemented with 100ng/mL of minimal epitope, 2ug/mL aCD28, 100IU/mL rhIL-2, and 2.5μM of CAL-101 or DMSO, as indicated. 1×10^6^ cells were plated per well in a 24-well plate and allowed to incubate at 37°C and 5% CO2. Cells were split after 48 hours by pooling cultured cells from all replicate wells, washing with PBS, counting, and re-plating at 1×10^6^ cells per well. rhIL-2, DMSO, and CAL-101 were replenished after splitting. Cells were incubated for another 48 hours before expanded CD8^+^ T cells were harvested, washed, resuspended in PBS, and counted. 10^5^ (OT-1) or 10^6^ (p-mel) CD8^+^ T cells were transferred into recipient mice by IV tail vein injection for ACT experiments as indicated.

### Vaccines and immunizations

SNP-7/8a vaccines were synthesized as previously described^45^. Peptides used in nanoparticles within the SNP-7/8a vaccine were ordered from GenScript. The hydrophobic blocks necessary for the SNP-7/8a vaccine were formulated by Vaccitech North America and linked to peptides using click chemistry. Formulated SNP-7/8a vaccines were validated by high-performance liquid chromatography and dissolved in 100ul sterile PBS for administration. Ova:SNP-7/8a, E7:SNP-7/8a, and Ova:SNP vaccines were administered IV or IM at 8nmol, as indicated. IV vaccines were administered by tail vein injection while IM vaccines were injected bilaterally (50ul on each side) into the gastrocnemius muscle. In all experiments using gp100:SNP-7/8a, vaccine was administered IV at 32nmol.

### Blood and tissue harvest

Unless otherwise indicated, whole blood was collected from animals 7d after ACT. Blood was collected in tubes containing heparin before removing red blood cells with ACK buffer. PBMCs were then washed in PBS and stained for flow cytometry. For organ harvest experiments, mice were euthanized 3 days following vaccination and spleens, tdLNs, and tumors were dissected. Spleens were mechanically dissociated and lysed with ACK buffer. tdLNs were mechanically dissociated with BioMasher tubes (Nippi). Tumors were placed in RPMI 1640 media containing 10% FCS, 50 U/ml DNase I (Sigma-Aldrich) and 0.2 mg/ml collagenase D (Sigma-Aldrich) and dissociated into single cell suspensions using gentleMACS dissociator (Miltenyi Biotec) per manufacturers instructions. All single cell suspensions were then washed in PBS, filtered through a 70 micron filter, and resuspended in PBS.

### Flow cytometry

For flow cytometry staining, cells were first stained with the LIVE/DEAD Fixable Blue Dead Cell Stain Kit (Invitrogen) in FACS buffer containing 2% FBS and 50nM dasatinib (Stemcell technologies) for 30 minutes at room temperature. For tetramer stain, cells were then washed, blocked with anti-CD16/CD32, and stained with the H2-Kb SIINFEKL tetramer (NIH Tetramer Core Facility) conjugated to streptavidin PE in FACS buffer. Simultaneously, cells underwent surface staining for 1 hour at 4°C with antibodies probing for: CD8 (clone 53-6.7), PD-1 (clone 29F.A12), CD62L (clone MEL-14), TIM-3 (clone RMT3-23), CD39 (clone Duha59), CD127 (clone A7R34) and KLRG1 (clone 2F1), CD45.2 (clone 104), and SLAMF6 (clone13G3) purchased from BioLegend and CD4 (clone RM4-4), CD44 (clone IM7), CD69 (clone H1.2F3), and CD45.1 (clone A20) purchased from BD Biosciences. Cells were then washed and fixation/permeabilization was performed using the Foxp3 Transcription Factor Staining Buffer kit (ebioscience) per manufacturers instructions. Fixed and permeabilized cells were then stained intracellularly overnight at 4°C with antibodies probing for: CD3 (clone 145-2C11) and KI-67 (clone B56) purchased from BD Biosciences and TCF1 (clone C63D9) from Cell Signaling Technology.

For intracellular cytokine staining of splenocytes following peptide stimulation, cells were first stained with the LIVE/DEAD Fixable Blue Dead Cell Stain Kit (Invitrogen) in FACS buffer containing 2% FBS for 10min at room temperature. Cells were then washed, blocked with anti-CD16/CD32, and stained with same surface antibodies for 20min at room temperature. Cells were then washed and fixation/permeabilization was performed using the BD Bioscienes Cytofix/Cytoperm kit. Cells were intracellularly probed for IFN-γ (clone XMG1.2).

Cells were analyzed by an LSRFortessa X50 (BD Biosciences) and all flow cytometry data was analyzed using FlowJo (Tree Star).

### IFN ELISA and Luminex

Serum was collected 6 hours following vaccination into serum separator tubes. ELISA kits were used to measure IFN-α and IFN-β (PBL Assay Science) and ran in accordance with manufacturer’s instructions. A broad serum cytokine panel was ran using the MILLIPLEX^®^ Mouse Cytokine/Chemokine Magnetic Bead Panel (Millipore Sigma) and ran in accordance with manufacturer’s instructions.

### Generation of scRNAseq libraries

For scRNAseq libraries from cultured cells, T_scm_ and T_eff_ cells were harvested at the end of the four day culture protocol and a total of 4×10^5^ cells were ran for each treatment group (1×10^5^ cells/lane). Single cell libraries were generated in accordance with manufacturer instructions for the Chromium Next GEM Single Cell 3ʹ v3.1 kit (10x Genomics). Sequencing was performed on a NovaSeq 6000 S1 chip (Illumina).

For scRNAseq libraries from mouse tumors, tumors were harvested and prepared as single cell suspensions 24 hours following vaccination. Tumor suspensions were first stained with the LIVE/DEAD Fixable Blue Dead Cell Stain Kit (Invitrogen) in FACS buffer containing 2% FBS for 30 minutes at room temperature. Samples were washed and stained with an antibody cocktail composed of anti-CD16/CD32 Fc block and CD45 APC (clone QA17A26). Tumor suspensions of individual treatment groups (T_eff_ cells, T_scm_ cells, T_eff_ cells + Vax, T_scm_ cells + Vax) were also incubated with a 1:1600 dilution of a hashtag antibody (TotalSeq anti-mouse Hashtag antibodies CMO301-CMO304, BioLegend). Cells were stained for 1 hour at 4°C, washed, and resuspended in FACS buffer before sorting. Live, CD45^+^ cells were sorted and pooled into the same tube. To tag cell surface proteins with oligo-conjugated antibodies, sorted cells were first blocked with anti-CD16/CD32 in FACS buffer for 10 minutes at 4°C. A commercially available 102-plex antibody cocktail (TotalSeq B Mouse Universal Cocktail V1, BioLegend) was then added to cells and incubated for 30 minutes at 4°C. Reaction was quenched in FACS buffer and cells were washed and resuspended in PBS. 9.8×10^4^ cells were loaded per lane and single cell gene expression (GEX) and cell surface protein (CSP) libraries were generated in accordance with the Chromium Next GEM Single Cell 3’ HT Kit v3.1. Sequencing was performed on a NovaSeq X 10B chip (Illumina).

### Analysis of tumor scRNAseq libraries

Raw output base call files were converted to FASTQ files and aligned to the mm10-3.0.0 mouse reference genome using the Cell Ranger v7.1.0 analysis pipeline (10x Genomics). FASTQ files were used to generate a count matrix composed of unique molecular identifiers using the Seurat R package v5.1.0^89^. Cells with low quality reads were removed and remaining cells were normalized and scaled in accordance with Seurat analysis guidelines. Dimensional reduction via principal component analysis (PCA) was performed and cells were clustered by the Louvain algorithm and projected via uniform manifold approximation and projection (UMAP). Clusters were annotated based on expression of DEGs, canonical markers, and CSP for tumor clusters. Volcano plots of DEGs were produced using the FindMarkers function (Seurat) and filtering results to include those with adjusted p values < 0.05 and log_2_ Fold Change values > 1. Gene set enrichment analysis (GSEA) was performed using the molecular signatures database hallmark gene set^90^. Gene ontology (GO) enrichment analysis was performed using Biological Process GO terms and the clusterProfiler R package^91,92^. RNA velocity analysis was performed using velocyto^93^ and scvelo^94^; pseudotime cell lineage analysis was performed using slingshot^95^. Dotplots and alluvial plots were made using ggplot2.

### Tumor fixation and generation of spatial transcriptomic libraries

Three days following vaccination, mouse tumors were harvested and washed in PBS. Tumors were submerged in 4% paraformaldehyde (PFA) and incubated for 24 hours at room temperature with gentle rocking. Following incubation, PFA was removed and tissue was washed three times with distilled water. Samples were stored in 70% ethanol at 4°C before paraffin embedding. Formalin-fixed, paraffin-embedded (FFPE) blocks were maintained at 4°C before being sectioned and stained with hematoxylin and eosin (H&E) by Histoserv (Germantown, MD). High resolution images of each tumor H&E were captured using a Leica SP5 microscope equipped with a Leica DFC425 C camera in collaboration with the Research Technologies Branch (NIAID/NIH). Samples were processed for the Visium HD assay in accordance with manufacturer instructions (CG000684 & CG000685, 10x Genomics). Briefly, H&E slides were destained and decrosslinked before probes were hybridized onto tissue RNA. Hybridized probe-RNA pairs are then ligated and within the Cytassist instrument are released from tissue, captured on Visium HD slide, and extended with a spatial barcode and UMI. Constructs were amplified and indexed for Illumina sequencing. Sequencing was performed on a NovaSeq X 1.5B chip (Illumina).

### Cell segmentation, barcode assignment, and clustering of spatial transcriptomic data

Cell segmentation was performed using Cellpose 2.0 by training custom models to identify cells within each TME image^96^. Following segmentation, cell masks were saved as polygon coordinates and imported into Rstudio to join barcode coordinates with polygon coordinates. Similar to previously published methods^97,98^ and using code adapted from 10x Genomics^99^, all 2×2μm barcodes contained within the perimeter of a given cell polygon were assigned to that cell and aggregated. A cell count matrix composed of segmented cells with corresponding transcript counts was generated using all cell polygons contained within an image. Cell matrices from each treatment group were merged and filtered for dead/dying cells. Cells were normalized and scaled in accordance with Seurat spatial analysis guidelines. PCA reduction, Louvain clustering, and UMAP were performed, resulting in clusters that were annotated based on differential gene expression and canonical markers. Images containing UMAP clustering information were projected using ggplot2.

### Spatial neighborhood analysis

Analysis of tumor-immune regions was performed by dividing images into tiled regions such that each image had ∼25 cells/tile in tiles containing at least two cells. Tiles containing at least one tumor cell (Stable tumor, Progressive tumor, or IFN-reactive tumor) and at least one immune cell (Stimulatory macs, Tumor resident macs, Proliferative stimulatory macs, Proliferative tumor macs, Monocytes, or T cells) were highlighted in red as a tumor-immune interaction tile using geom_tile function of ggplot2. Density of tumor and immune cells were calculated by fractioning the number of tumor or immune cells in a tile by the total number of cells in that tile. Statistics on immune cell densities were calculated by extracting data from all tiles containing at least one relevant immune cell and fractioning the number of relevant immune cell by the total number of cells in each tile and performing a two-sided Wilcoxon test. Z scores were calculated using the neighborhood enrichment function in Squidpy^100^. Spatial ensembles and patterns were calculated and plotted with the SPACE^59^ package using a 5x tissue coverage and observing differences across a radius distance of 60μm with the measure_cisMI function.

### Pathway enrichment and DEG analysis from spatial transcriptomic data

The FindMarkers function (Seurat) was used to create a ranked gene list between the TMEs of mice treated with T_scm_ cells + Vax and T_eff_ cells + Vax. GSEA was performed using the molecular signatures database hallmark gene set and GSEA enrichment plots were made using the enrichplot R package. All DEGs (p < 0.05 and log_2_FC > 1) were calculated using the FindMarkers function. DEGs between T_scm_ cells + Vax and T_eff_ cells + Vax were annotated based on biological function and graphed as a heatmap using the pheatmap R package. Overlapping DEGs between treatment groups were graphed as a Venn diagram using the ggvenn R package. DEGs between T_scm_ cells alone and T_scm_ cells + Vax were converted to human homologues using the homologene R package to create a list of huDEGs. The TME DEGs of responders vs non-responders to ACT were obtained from Lowery et al. 2025 and overlapped with huDEGs, resulting in 71 matched genes. Matched genes from both studies were plotted by log_2_FC expression and were termed “Positively concordant” if their log_2_FC expression was positive in both studies, and “Negatively concordant” if their log_2_FC expression was negative in both studies. A Pearson correlation test was used to calculate the correlation coefficient (r) and p value. Enrichment plots were created by calculating the running enrichment score of ranked huDEGs against genes upregulated in responders to ACT or upregulated in non-responders to ACT.

### TCGA analysis

TCGA clinical data was obtained from the Genomic Data Commons (NIH/NCI) and filtered to only include primary tumors (sort code 01). Matched, batch corrected, TCGA pan-cancer mRNA data (n = 11,060) were obtained from the UCSC Xenabrowser (https://xenabrowser.net) and gene-wise Z scoring was performed to achieve equal weighting of genes. Samples with NA expression values or without survival data were removed from analysis. A signature score was defined for each patient by calculating the difference between the average Z scored expression of huDEG-upregulated genes and huDEG-downregulated genes. Patients (n = 8077) were stratified into high or low expression groups based on the median signature score. Kaplan-Meier survival curves were generated in R using the survival package and p values were calculated using a log-rank test.

### Statistics

Statistics were accessed by a paired, two-tailed T test (cultured T cell phenotyping), two-way ANOVA with Bonferroni’s correction (tumor protection curves), one-way ANOVA with Tukey post-hoc test (multiple comparisons by flow cytometry phenotyping and cytokine assays), log-rank test (survival analyses), and two-sided Wilcoxon test (cell density across TMEs). Statistics were performed in Rstudio or Prism (GraphPad). P values with asterisks are reported as P < 0.05 (*), P < 0.01 (**), P < 0.001 (***), P < 0.0001 (****).

## Acknowledgments

We thank all members of the Seder and Cortés-Ciriano labs for helpful discussion and input. We also thank members of the Translational Research Program (Vaccine Research Center) for their critical help with animal studies, especially Terri Bennaugh, Nina Callaham, Marqui Dorsey, Sienna Rush, Lesly Mendoza, Hilda Quintanilla, Reina Quintanilla, and Carmelo Chiedi. We are grateful to the VRC Genome Analysis Core, particularly Amy Ransier and Farida Laboune for their help with sequencing. We thank the Research Technologies Branch (NIH/NIAID), including Owen M. Schwartz and Margery Smelkinson for imaging help. We are also grateful to members of Barinthus Biotherapeutics for support with the manufacture of SNP-7/8a vaccines. This research was supported by the Intramural Research Program of the National Institutes of Health (NIH). The contributions of the NIH authors are considered Works of the United States Government. The findings and conclusions presented in this paper are those of the authors and do not necessarily reflect the views of the NIH or the U.S. Department of Health and Human Services.

