## Supplementary Figures S1-S10 for "Spatiotemporal dynamics of adoptively transferred stem-like CD8^+^ T cells in the tumor microenvironment following vaccination"

### Figure S1

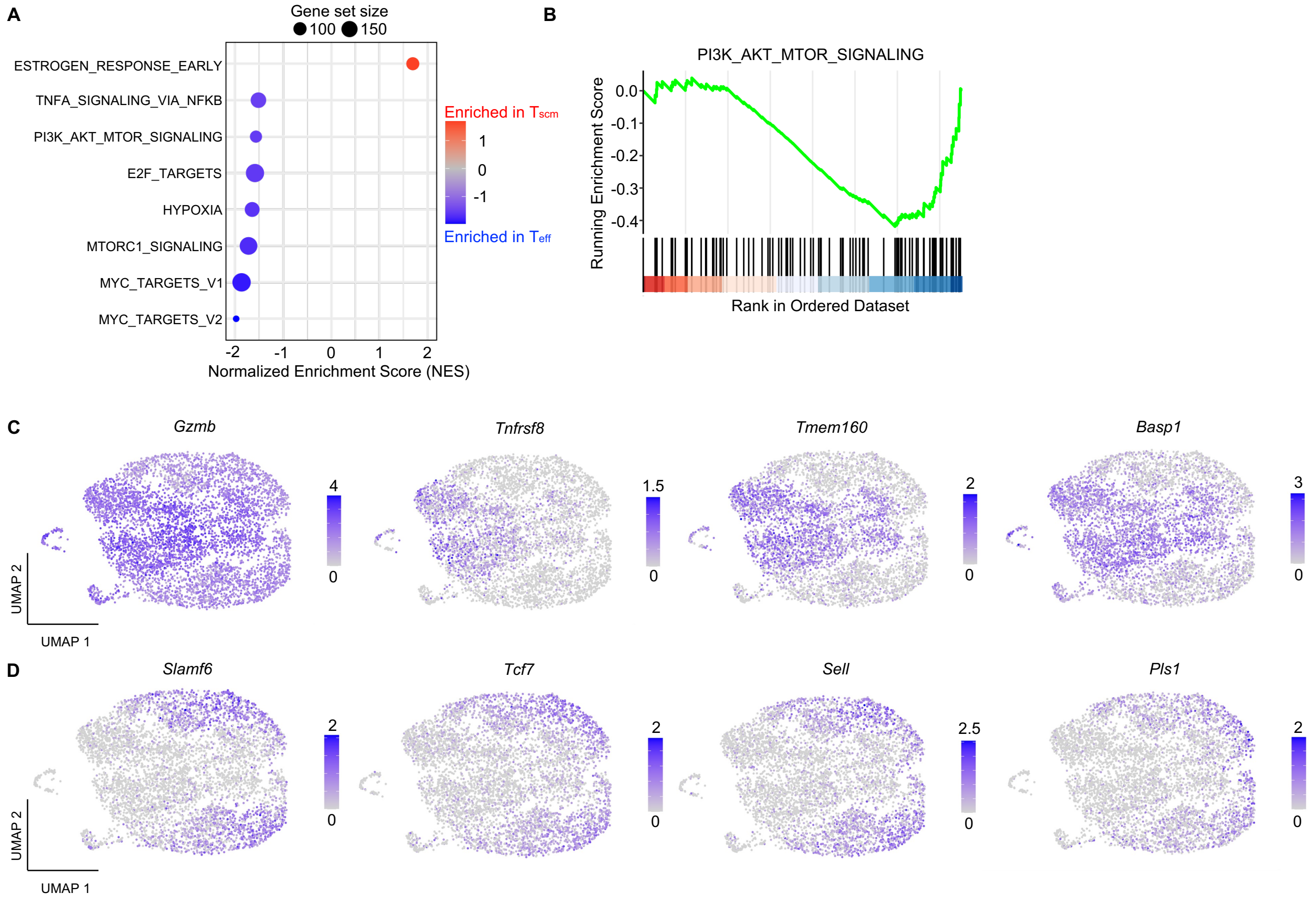


**Figure S1. Analysis of global gene expression patterns in $T_{scm}$ cells or $T_{eff}$ cells by scRNAseq**
(A) Dot plot indicating Hallmark gene pathways enriched in $T_{scm}$ cells or $T_{eff}$ cells.
(B) Enrichment plot showing running enrichment score across ranked DEGs between $T_{scm}$ cells and $T_{eff}$ cells for the PI3k/AKT/mTOR signaling pathway.
(C) Feature plots indicating expression of DEGs related to $T_{eff}$ cells.
(D) Feature plots indicating expression of DEGs related to $T_{scm}$ cells.

#### Figure S2

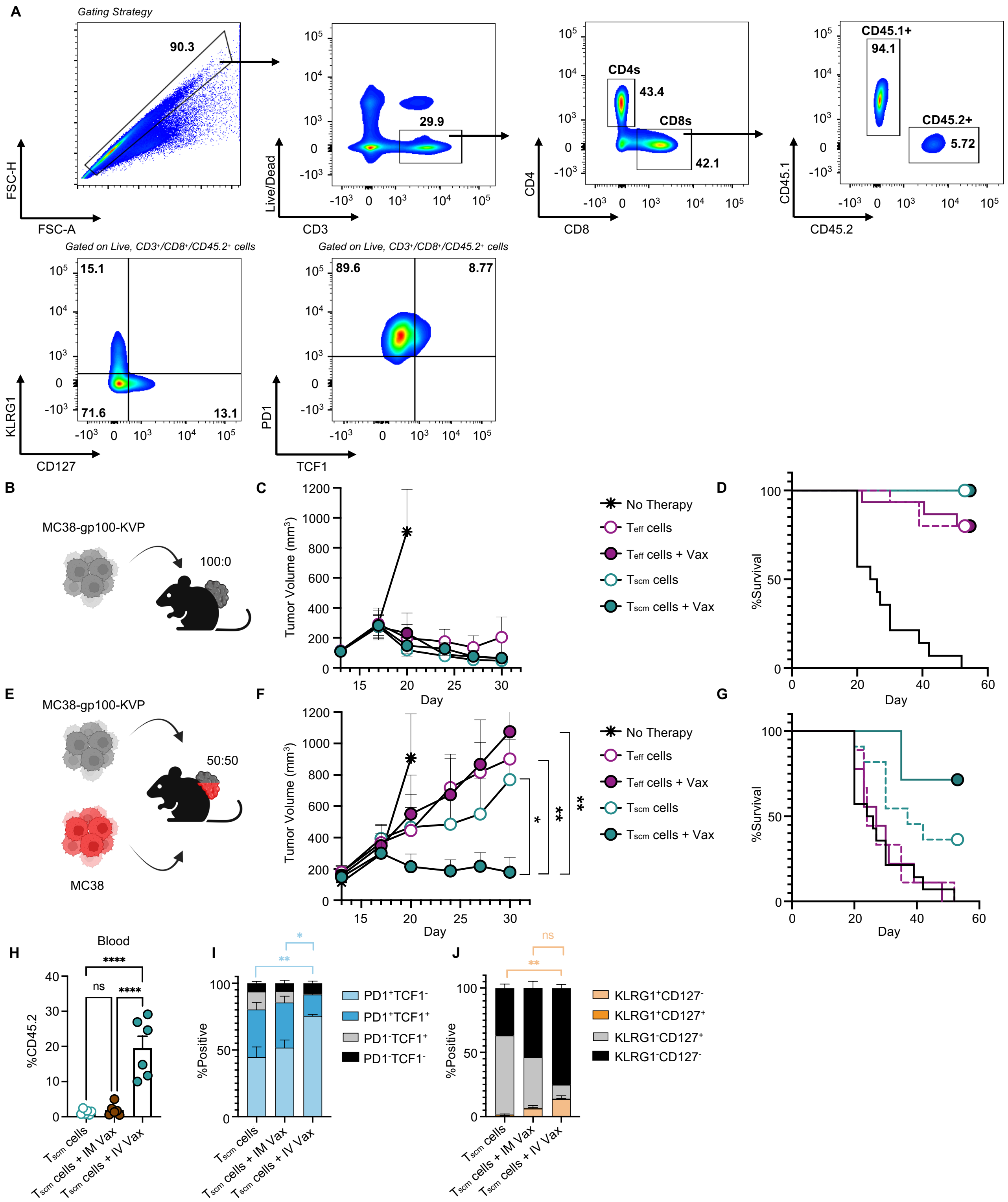


**Figure S2. Effect of ACT and vaccination in an MC38 tumor model**

(A) Flow cytometry gating strategy for CD8+ T cell phenotyping of tumor, spleen, and tdLN cell suspensions.

(B) Experimental schematic of mouse model testing ACT and vaccination on mice inoculated with 100% MC38-gp100-KVP cells.

(C) Tumor curve following treatment of $T_{eff}$ or $T_{scm}$ cells with or without vaccination with gp100:SNP-7/8a (n=10) in 100% MC38-gp100-KVP tumor setting. Statistics accessed by two-way ANOVA.

(D) Survival curve following treatment of $T_{eff}$ or $T_{scm}$ cells with or without vaccination with gp100:SNP-7/8a (n=10) in 100% MC38-gp100-KVP tumor setting.
(E) Experimental schematic of mouse model testing ACT and vaccination on mice inoculated with a 50:50 mixture of MC38-gp100-KVP and WT MC38 cells.
(F) Tumor curve following treatment of $T_{eff}$ or $T_{scm}$ cells with or without vaccination with gp100:SNP-7/8a (n=10) in 50:50 MC38-gp100-KVP:MC38 tumor setting. Statistics accessed by two-way ANOVA.
(G) Survival curve following treatment of $T_{eff}$ or $T_{scm}$ cells with or without vaccination with gp100:SNP-7/8a (n=10) in 50:50 MC38-gp100-KVP:MC38 tumor setting
(H) Bar graph showing the frequency of CD45.2$^{+}$ transferred cells in the blood following ACT and IM or IV vaccination (n=6). Statistics accessed by ANOVA.
(I) Stacked bar graphs indicating the percentages of PD1 and TCF1 subpopulations in blood (n=6). Statistics assessed by ANOVA.
(J) Stacked bar graphs indicating the percentages of KLRG1 and CD127 subpopulations in blood (n=6). Statistics assessed by ANOVA.

### Figure S3

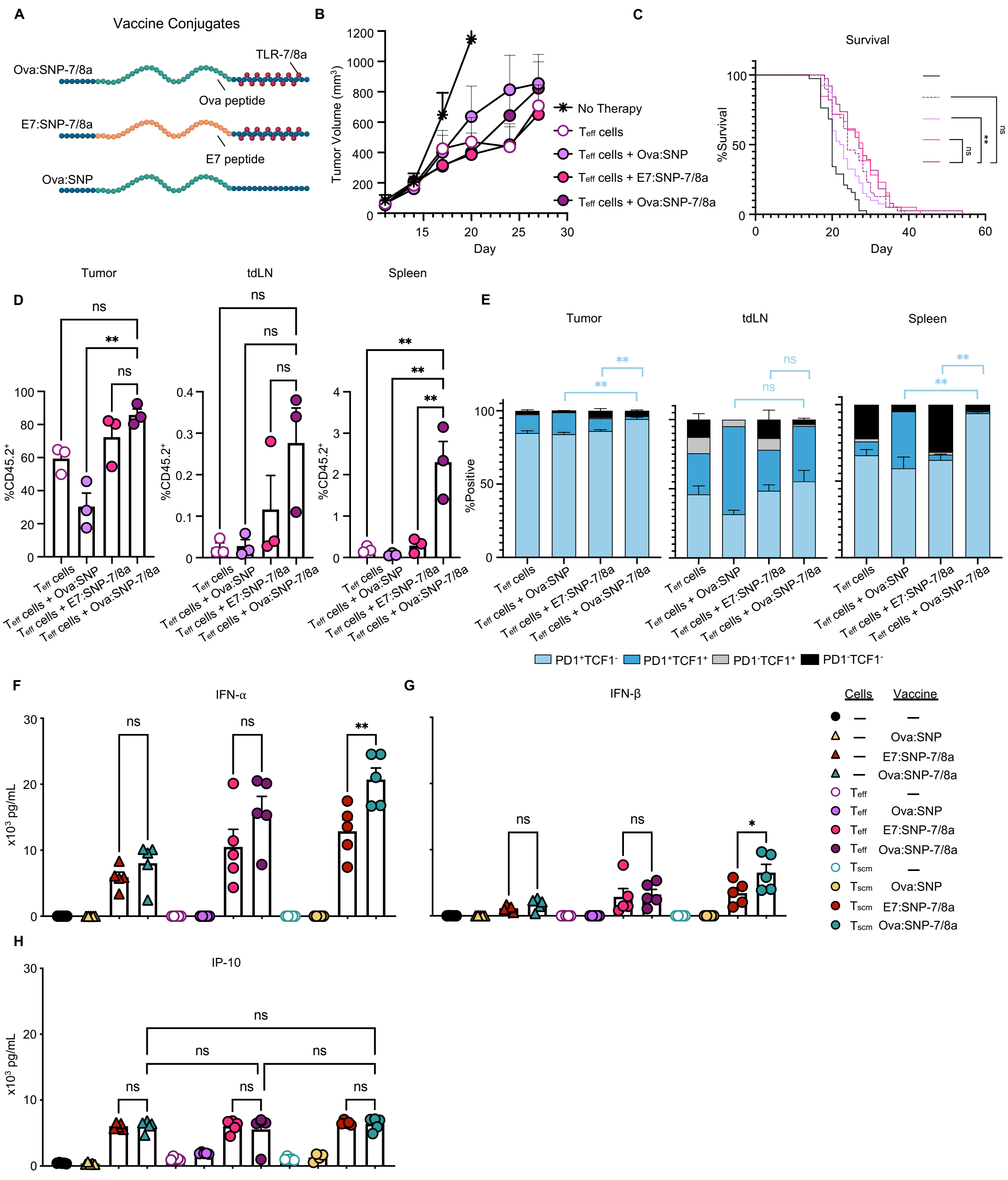


**Figure S3. Effect of antigen and innate stimulation following ACT with $T_{eff}$ cells**

(A) Cartoon detailing the vaccine conjugates used in experiments: Ova:SNP-7/8a (innate + antigen), E7:SNP-7/8a (innate alone), Ova:SNP (antigen alone).
(B) Tumor curve following treatment of $T_{eff}$ cells with various vaccine conjugates (n=7). Statistics accessed by two-way ANOVA.
(C) Survival curve following treatment of $T_{eff}$ cells with various vaccine conjugates (n=35). Statistics accessed by log-rank test.
(D) Bar graphs showing the frequency of $CD45.2^+$ transferred cells in the tumor, tdLN, and spleen by treatment group (n=3). Statistics accessed by ANOVA.
(E) Stacked bar graphs indicating the percentages of PD1 and TCF1 subpopulations in tumor, tdLN, and spleen (n=3). Statistics assessed by ANOVA.
(F) Bar graph showing the serum cytokine levels (pg/mL) of IFN-α 6 hours post-vaccination in indicated treatment groups (n=5). Statistics assessed by ANOVA.
(G) Bar graph showing the serum cytokine levels (pg/mL) of IFN-β 6 hours post-vaccination in indicated treatment groups (n=5). Statistics assessed by ANOVA.
(H) Bar graph showing the serum cytokine levels (pg/mL) of IP-10 6 hours post-vaccination in indicated treatment groups (n=5). Statistics assessed by ANOVA

#### Figure S4

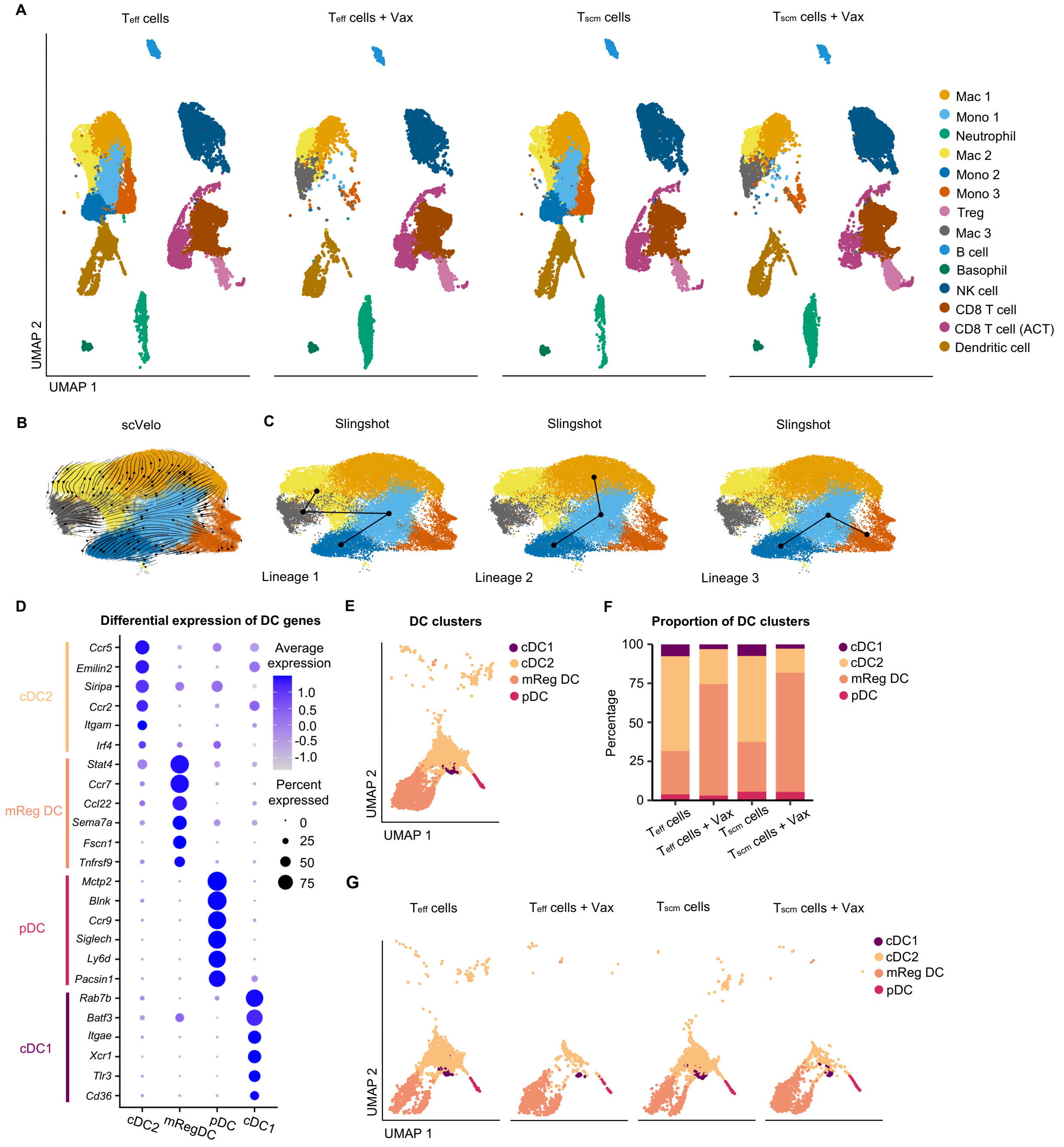


**Figure S4. Cell trajectory analysis of MonoMac clusters and dendritic cell subclustering in scRNAseq of TME**
(A) UMAP plots showing all cell clusters present in indicated treatment groups 24 hours following vaccination.
(B) scVelo analysis plotted over monocyte and macrophage cell clusters combined from each treatment group. Arrows indicate RNA velocity in cells throughout UMAP.
(C) Cell trajectory plots showing possible cell lineages derived from Slingshot analysis. Each lineage begins at the Mono 2 cluster and lines indicate cellular trajectories.
(D) Dot plot highlighting differentially expressed genes used to define and annotate subclustered dendritic cell populations. Average and percent expression is indicated by size and color of dots.
(E) UMAP plot showing the subclusters of dendritic cells combined from all treatment groups.
(F) Stacked bar graph showing the proportion of dendritic cell subclusters across indicated treatment groups.
(G) UMAP plots showing the dendritic cell subclusters present in indicated treatment groups 24 hours following vaccination.

### Figure S5

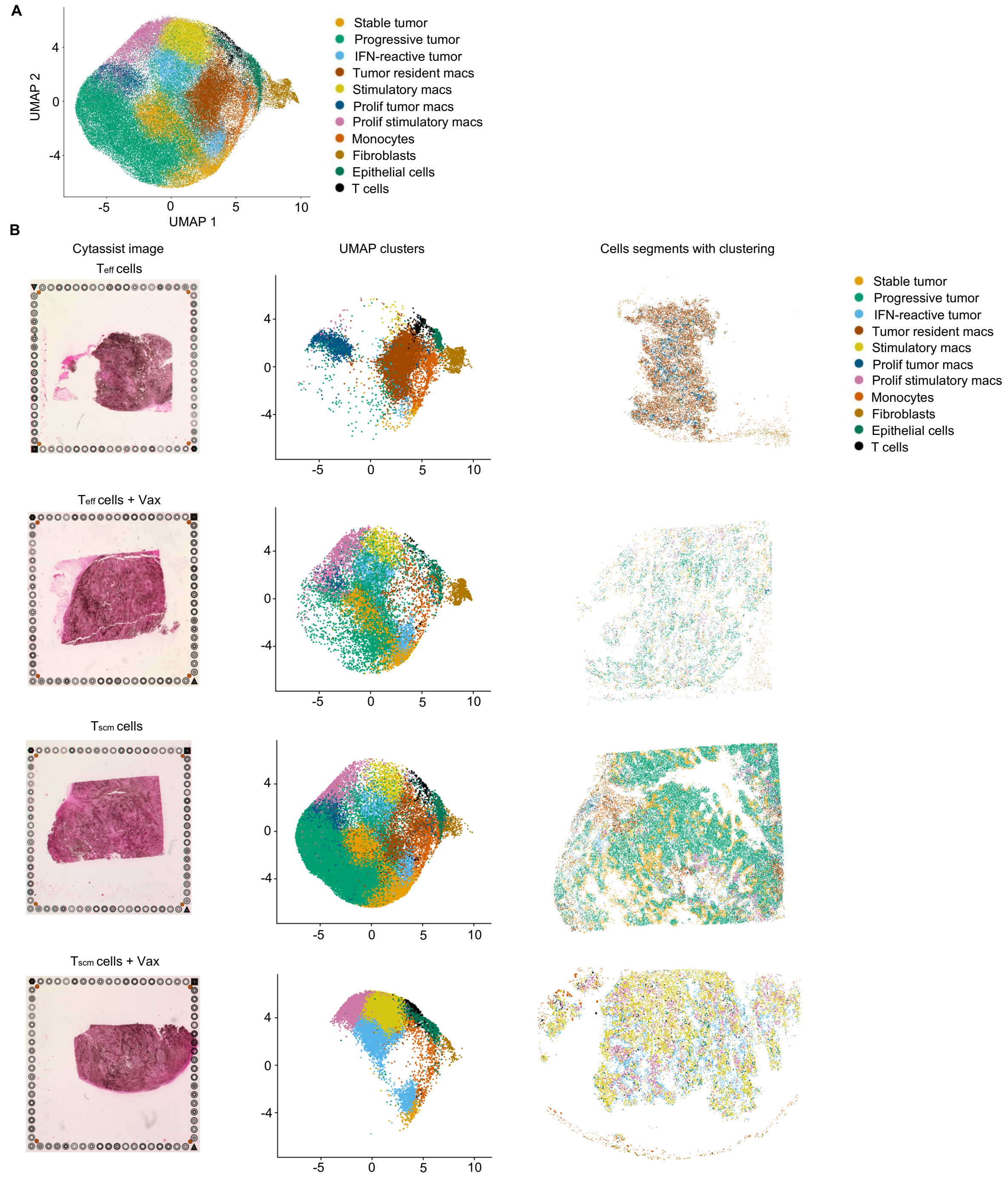


**Figure S5. Cell type annotation from spatial transcriptomics data**
(A) UMAP from spatial transcriptomics data detailing annotated clusters of cells present in the TME three days following vaccination.
(B) Panels showing the raw cytassist image (left), cell clusters present in UMAP (middle), and spatial feature plot of all segmented cells (right) in indicated treatment group.

### Figure S6

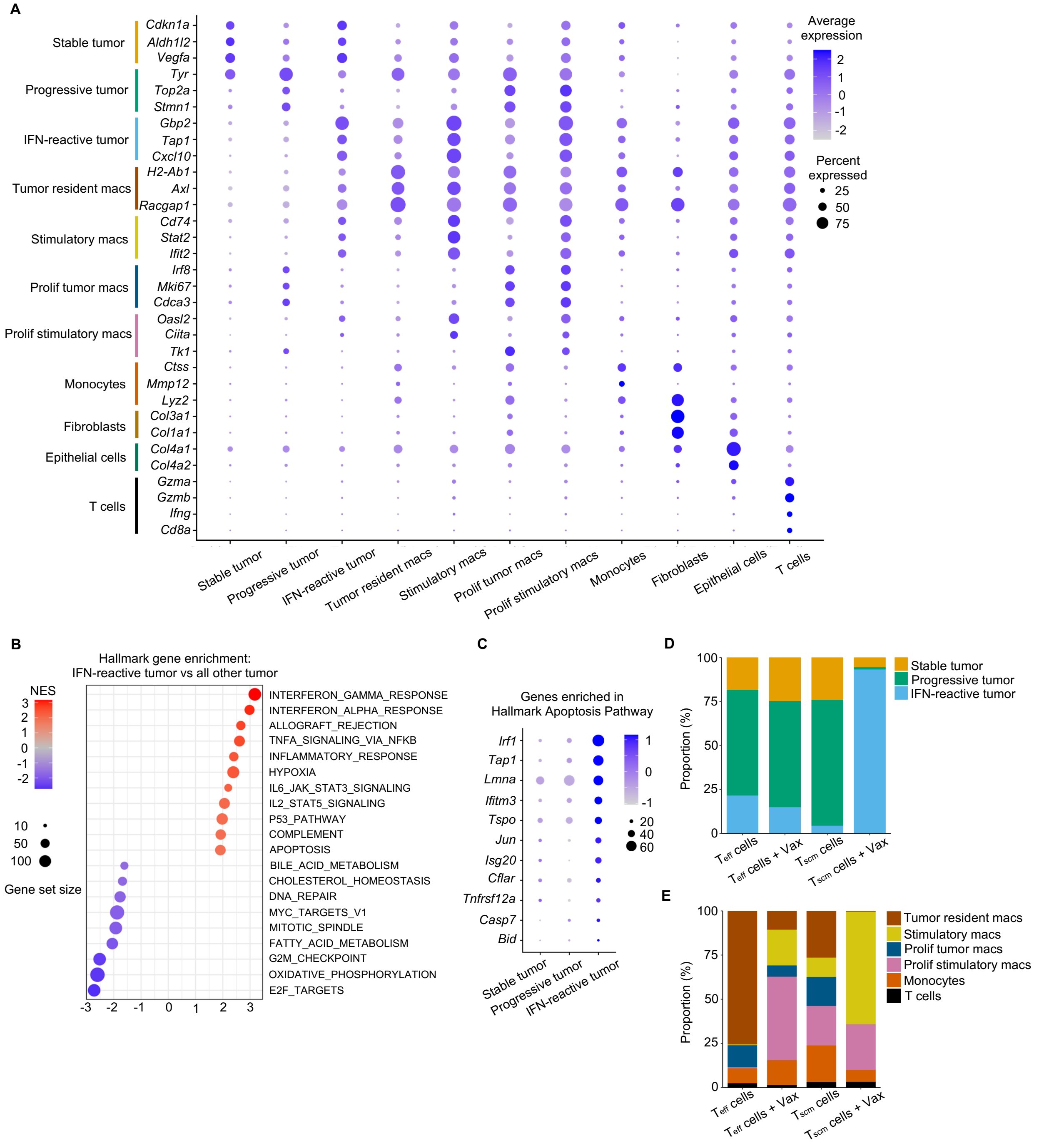


**Figure S6. Spatial description of treatment groups by tumor and immune clusters**

(A) Dot plot highlighting differentially expressed genes used to define and annotate each cluster present in the TME of spatial transcriptomics data.

(B) Dot plot indicating Hallmark gene pathways upregulated (red) or downregulated (blue) in IFN-reactive tumor cells compared to Stable and Progressive tumor cells.

(C) Dot plot highlighting genes within the Hallmark_APOPTOSIS pathway that are enriched in IFN-reactive tumor cells compared to Stable and Progressive tumor cells.

(D) Stacked bar graph showing the proportion of tumor cell clusters across indicated treatment groups.

(E) Stacked bar graph showing the proportion of immune cell clusters across indicated treatment groups.

#### Figure S7

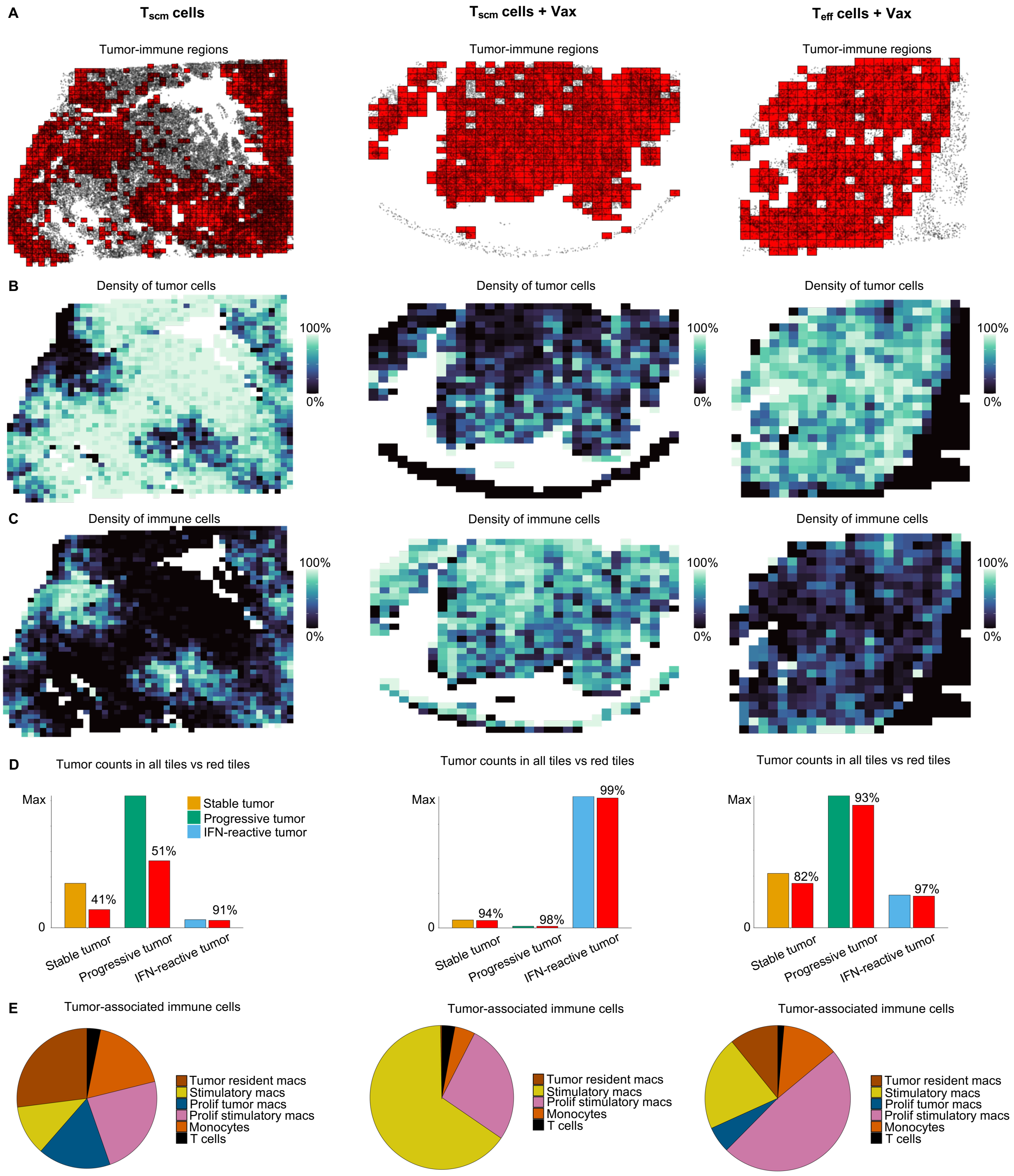


**Figure S7. Effect of intravenous vaccination on an immune-active TME**
(A) Spatial feature plots of indicated treatment groups showing the presence of tumor-immune interactions in local tiled neighborhoods (red).
(B) Spatial feature plots of indicated treatment groups displaying a scaled heatmap of tumor cell density within each tiled neighborhood.
(C) Spatial feature plots of indicated treatment groups displaying a scaled heatmap of immune cell density within each tiled neighborhood.
(D) Grouped bar graphs of indicated treatment groups showing total counts of each tumor type next to counts of each tumor type only contained within red tiles.

(E) Pie charts of indicated treatment groups showing the proportion of immune cells contained within the red tiled neighborhoods of each TME.

#### Figure S8

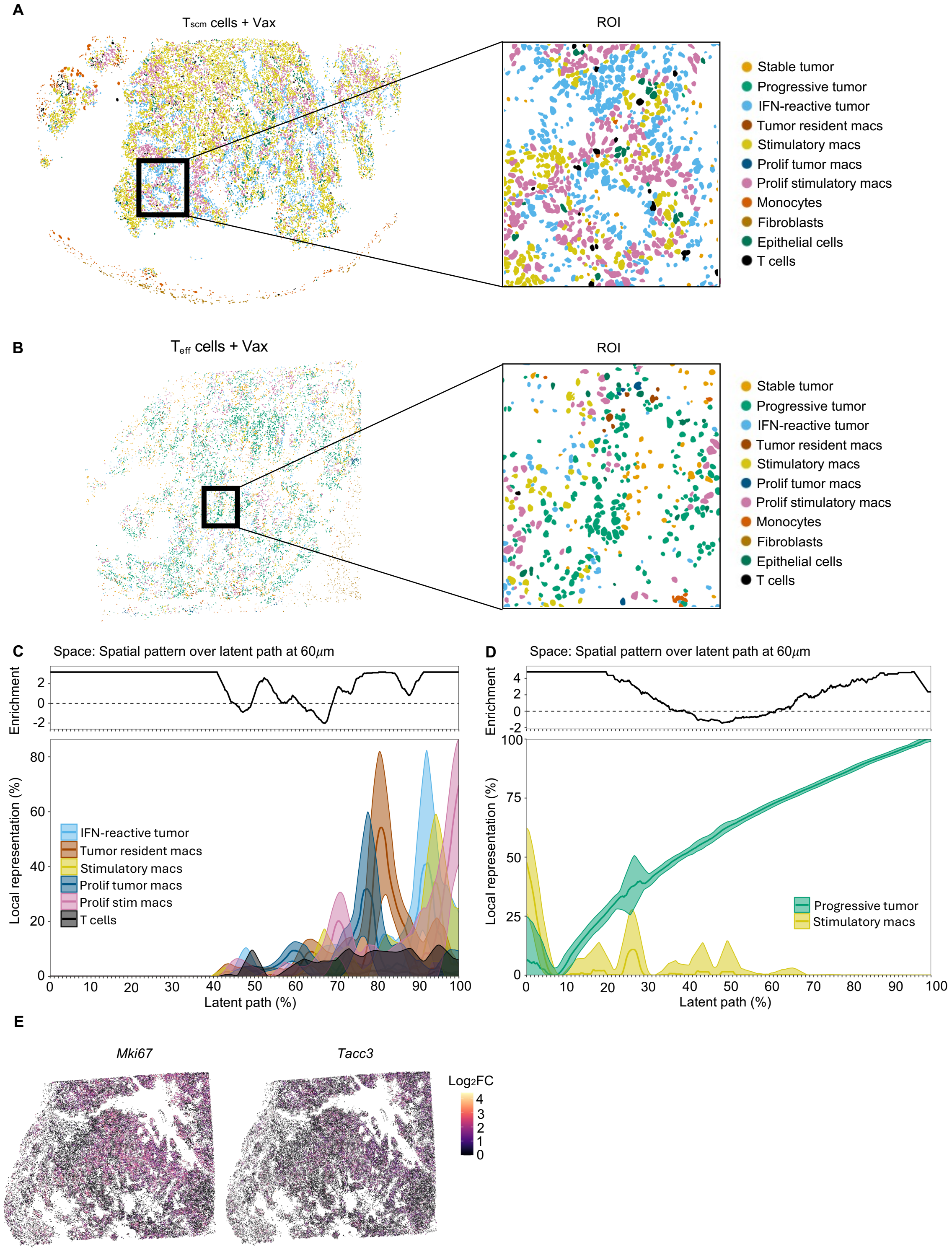


**Figure S8. Investigation of tumor-immune cell interactions by spatial neighborhood analysis**

(A) Spatial feature plot of segmented cells in TME treated with $T_{scm}$ cells + Vax. Region of interest highlights a zoomed-in, representative image of the TME.

(B) Spatial feature plot of segmented cells in TME treated with $T_{eff}$ cells + Vax. Region of interest highlights a zoomed-in, representative image of the TME.

(C) Covariation plot detailing the smoothed mean and 95% confidence interval of the local representation of all immune cells across latent path with running enrichment score in TME treated with $T_{scm}$ cells.

(D) Covariation plot detailing the smoothed mean and 95% confidence interval of the local representation of indicated cell types across latent path with running enrichment score in TME treated with $T_{scm}$ cells.
(E) Spatial feature plots of TME treated with $T_{scm}$ cells highlighting the $Log_2FC$ expression of *Mki67* and *Tacc3* within each cell.

#### Figure S9

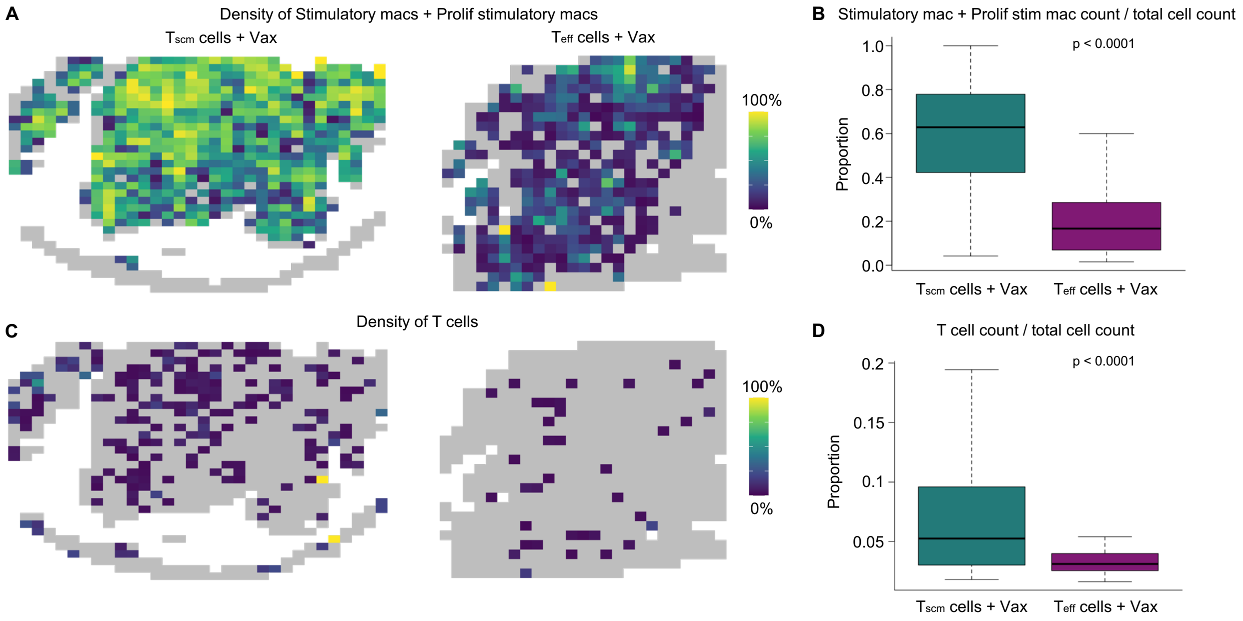


**Figure S9. Differences in immune infiltration in TME following treatment with $T_{scm}$ cells + Vax or $T_{eff}$ cells + Vax**
(A) Spatial feature plots of indicated treatment groups displaying a scaled heatmap of combined stimulatory macrophage and proliferative stimulatory macrophage cell density within each tiled neighborhood.
(B) Bar graph quantitating the proportion of combined stimulatory macrophages and proliferative stimulatory macrophages per tiled neighborhood in tiles containing at least one stimulatory macrophage in indicated treatment groups. Statistics accessed by a two-sided Wilcoxon test.
(C) Spatial feature plots of indicated treatment groups displaying a scaled heatmap of T cell density within each tiled neighborhood.
(D) Bar graph quantitating the proportion of T cells per tiled neighborhood in tiles containing at least one T cell in indicated treatment groups. Statistics accessed by a two-sided Wilcoxon test.

#### Figure S10

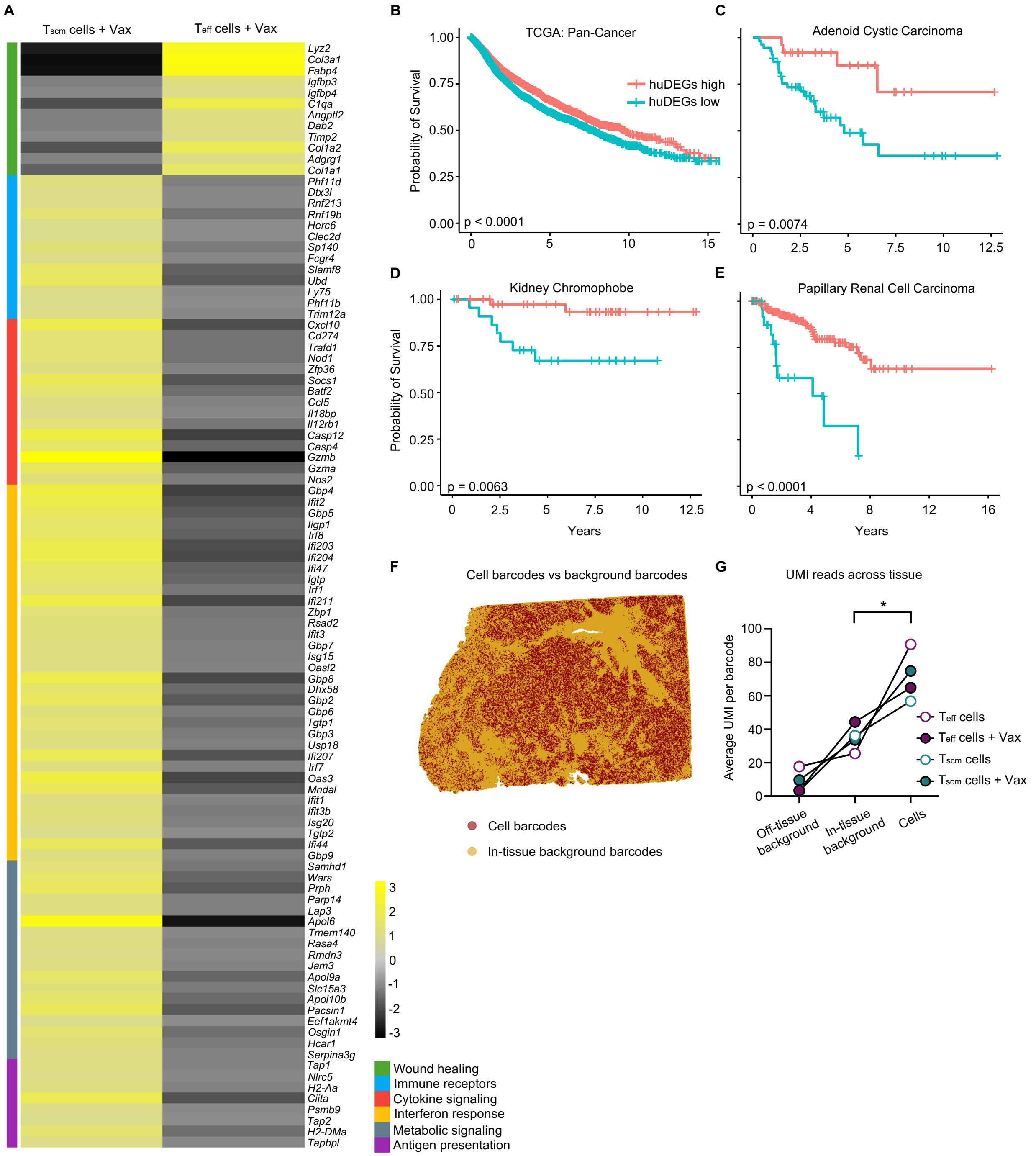


**Figure S10. Transcriptomic TME signatures in mice and humans with improved outcomes**

(A) Heatmap highlighting DEGs between TMEs treated with $T_{scm}$ cells + Vax and $T_{eff}$ cells + Vax. Plot is scaled based on $log_2FC$ expression and genes are annotated based on biological function.

(B) Kaplan-Meier survival curves from TCGA patient data across all tumors (pan-cancer). Patient cohorts were stratified as high (red) or low (blue) based on huDEGs signature score. Statistics accessed by log-rank test.

(C) Kaplan-Meier survival curves from TCGA patient data of Adenoid Cystic Carcinoma. Patient cohorts were stratified as high (red) or low (blue) based on huDEGs signature score. Statistics accessed by log-rank test.

(D) Kaplan-Meier survival curves from TCGA patient data of Kidney Chromophobe. Patient cohorts were stratified as high (red) or low (blue) based on huDEGs signature score. Statistics accessed by log-rank test.

(E) Kaplan-Meier survival curves from TCGA patient data of Papillary Renal Cell Carcinoma. Patient cohorts were stratified as high (red) or low (blue) based on huDEGs signature score. Statistics accessed by log-rank test.

(F) Spatial feature plot highlighting the presence of UMI barcodes found within segmented cells (red) or within the tissue are outside segmented cells (yellow).

(G) Line graph showing the average UMI reads per barcode in areas outside the tissue (off-tissue background), within the tissue and outside segmented cells (in-tissue background), and within segmented cells (cells). Statistics accessed by two-way ANOVA.
